# A Tonic Signaling Code Predicts CAR-T Cell Efficacy in Diffuse Midline Glioma

**DOI:** 10.1101/2025.09.29.679095

**Authors:** Emily B. Deng, Xiaowen Zhong, Dazhuan Xin, Upendra K. Soni, Wenkun Ma, Xiao Huang, Mingjun Cai, Po-Yu Liang, Jun Bai, Qian Qin, Shreya Mishra, Ming Hu, Arman E. Bayat, Jiajie Diao, Mei Xin, Natasha Pillay-Smiley, Trent R. Hummel, Charles B. Stevenson, Jessica B. Foster, Peter de Blank, Scott Raskin, Carl Koschmann, Jose A. Cancelas, Yi Zheng, Q. Richard Lu

**Author notes:** These authors contributed equally to this work (Emily B. Deng, Xiaowen Zhong, Dazhuan Xin). **Correspondence:** Dr. Richard Lu, Cincinnati Children’s Hospital Medical Center, Cincinnati, OH 45229, USA.

## Abstract

Diffuse midline glioma (DIPG/DMG) is a uniformly fatal pediatric brain tumor with no effective cure. Although CAR T-cell therapy shows promise, clinical outcomes remain inconsistent due to limited persistence and premature exhaustion. Reliable predictive biomarkers are lacking, and proposed exhaustion or stemness markers provide limited utility. Here, we systematically compare multiple CAR-T constructs targeting clinically-relevant antigen B7-H3 and identify antigen-independent CAR activation, or tonic signaling, as a key determinant of therapeutic performance. We find that B7-H3 CAR-T cells with restrained tonic signaling display superior tumor killing, persistence, and resistance to exhaustion, along with reduced CAR membrane clustering, in patient-derived DIPG models. Integrated multi-omics and single-cell profiling further reveal a CAR-T tonic signaling–associated gene signature that outperforms conventional exhaustion or stemness markers in predicting therapeutic efficacy across multiple clinical trials, including DIPG and other tumor types. Together, these findings define a mechanistic and predictive framework to guide CAR design and improve clinical outcomes.

## INTRODUCTION

Diffuse intrinsic pontine glioma (DIPG), recently classified under the broader category of diffuse midline glioma (DMG), is one of the most lethal pediatric high-grade brain tumors with a median survival of less than one year despite intensive treatment ^1-3^. The infiltrative nature of DMG/DIPG, its critical location in the pons, and its intrinsic resistance to conventional therapies make it exceptionally difficult to treat ^1-3^.

Chimeric antigen receptor (CAR) T cell therapy has revolutionized the treatment paradigm for brain tumors including DMG/DIPG ^4-7^. However, the clinical outcomes of these treatments are inconsistent and mixed. A major barrier in CAR-T therapies lies in the difficulties to achieve sustainable tumor control, partially due to the inadequate potency and persistence of CAR-T cells in eradicating tumor cells ^8-12^. Given the profound heterogeneity of brain tumors, the physical and structural barriers to therapy, and the unique challenges of achieving efficacy and long-term CAR-T persistence within the CNS, identifying robust biomarkers capable of predicting CAR-T cell efficacy is critical for improving outcomes and achieving durable responses in CAR-T therapy for brain tumors such as DMG/DIPG.

T-cell exhaustion or stemness properties have been proposed as indicators of CAR-T therapy efficacy ^13-15^. However, reliance on these downstream events such as T cell exhaustion to assess CAR-T performance has shown limited predictive value ^14,15^. Currently, reliable biomarkers to predict therapeutic efficacy are lacking. Antigen-independent activation, also known as tonic signaling, is a major contributor to premature T cell differentiation and exhaustion during CAR-T cell therapy ^16-18^. Variations in single-chain variable fragment (scFv) epitope binding specificity and affinity can alter tonic signaling propensity, thereby affecting the therapeutic efficacy and persistence of CAR-T cells ^19-21^. Although previous studies have highlighted the impact of tonic signaling on T cell exhaustion, the underlying mechanisms remain poorly understood. The extent to which tonic signaling quantitatively shapes CAR-T cell fate and function, and how these effects correlate with patient outcomes, is still unclear. In particular, the mechanistic connections between tonic signaling and the transcriptional and epigenetic programs that govern CAR-T durability are incompletely defined—especially in solid tumors such as DMG/DIPG, where persistence remains a critical barrier.

To address these mechanistic gaps, we engineered a panel of CARs targeting the clinically relevant tumor antigen B7-H3 (CD276) and systematically dissected the effects of tonic signaling. B7-H3, a member of the B7 family of immune modulators, is broadly expressed in DMG/DIPG and other pediatric high-grade gliomas while being largely absent from normal brain, making it an attractive CAR-T target ^22-26^. By comparing these B7-H3–targeted CAR constructs currently in clinical trials ^23,27^, we identify a specific B7-H3 CAR-construct with restrained tonic signaling, which outperforms counterparts with higher tonic activity across multiple functional assays *in vitro* and *in vivo*. Integrated transcriptomic, epigenomic, and single-cell profiling reveal that restrained tonic signaling imprints gene regulatory programs that preserve CAR-T stemness and function. Importantly, we define a tonic signaling–associated gene signature that consistently outperforms conventional exhaustion or stemness markers in predicting CAR-T efficacy across independent clinical datasets, including DIPG/DMG and other tumor types. Together, our study establishes a mechanistic and predictive framework beyond the previous recognition that tonic signaling influences CAR-T activity. This framework not only reveals how tonic signaling imprints transcriptional and epigenetic programs in CAR-T cells but also identifies a potential clinically relevant biomarker that can inform therapeutic efficacy in DMG/DIPG and other solid tumors.

## RESULTS

### Identification of tonic signaling-subdued B7H3 CAR-T cells with enhanced functionality against DIPG

To systematically evaluate the levels of antigen-independent tonic signaling of different B7-H3 CARs under clinical investigation ^23,27^, we generated lentiviral vectors encoding human codon-optimized 376.96 (B7H3.BC), MGA271, and Hu8H9 CAR constructs (Figure 1A). CAR-T cells were engineered to express each construct, which included a signal peptide, scFv derived from the anti-B7-H3 monoclonal antibodies 376.96, MGA271, or Hu8H9, a CD8α hinge and transmembrane domain (CD8TM), a 4-1BB intracellular co-stimulatory domain, and a CD3ζ intracellular signaling domain.

**Figure 1.**
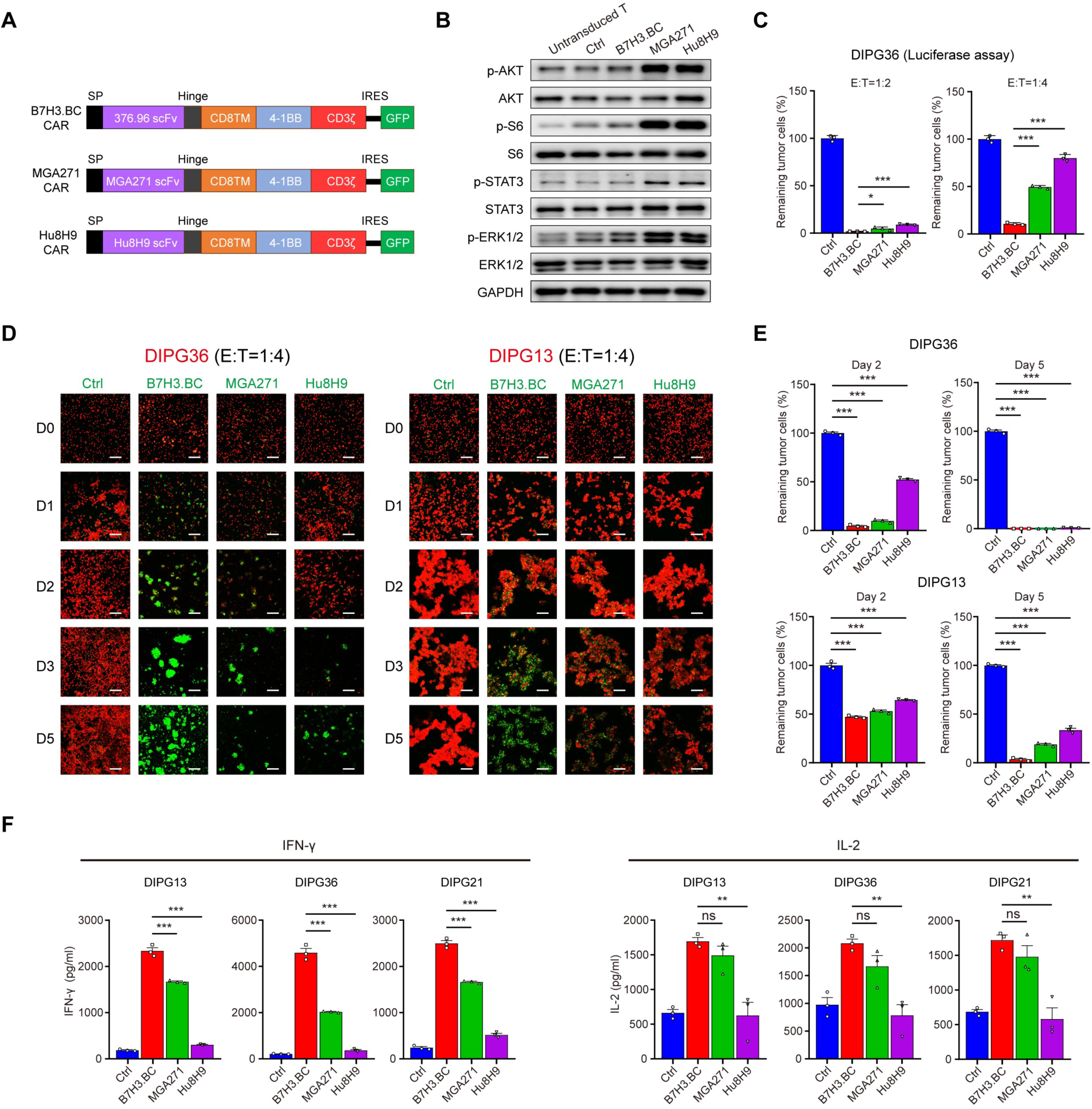
Identification of a B7H3 CAR with subdued tonic signaling and enhanced function against DIPG cells. **(A)** Schematic representations of lentiviral vectors encoding B7H3.BC, MGA271, and Hu8H9 CAR constructs. SP, signal peptide; Hinge, hinge region of human CD8α; CD8TM, CD8 transmembrane region; IRES, internal ribosome entry site. **(B)** Western blot analysis for phosphorylated and total S6, AKT, STAT3, ERK1/2 in B7-H3 CAR-T cells, T cells transduced with empty vector (CTRL), and untransduced T cells on day 10 of culture. Representative of three donors. **(C)** Percent survival of DIPG36 cells incubated with CAR-T lines relative to sample incubated with CTRL T cells assessed by luciferase assays after 2 days of co-culture. CAR-T effector to tumor cell ratios of 1:2 and 1:4. Data are means ± SEM from triplicate wells. Representative of three donors. **(D)** Fluorescence images T cells in co-culture with DIPG36 (left) and DIPG13 (right) cells. The ratio of T cells (green) to DIPG cells (red) was 1:4. Images were captured at days 0, 1, 2, 3, and 5. Scale bars = 200 µm. Representative of three donors. **(E)** Percentage survival of DIPG36 (upper) and DIPG13 (lower) at days 2 and 5 of incubation with indicated T cells. Data are means ± SEM. **(F)** Quantification of IFNγ and IL-2 release from T cells after 24 hours of co-culture with DIPG cells. Data are means ± SEM from triplicate wells. Representative of three donors. For panels **C**, **E** and **F**, unpaired two-tailed Student’s t-test. *P < 0.05, **P < 0.01, ***P < 0.001; ns, not significant.

To assess tonic signaling levels in these CAR-T cells, we measured phosphorylation of key signaling molecules previously associated with tonic signaling ^16,17,19,20^, including S6 (p-S6), AKT (p-AKT), STAT3 (p-STAT3), and ERK1/2 (p-ERK1/2). B7H3.BC CAR-T cells exhibited markedly lower basal phosphorylation of all four molecules compared with MGA271 and Hu8H9 CAR-T cells (Figure 1B), indicating reduced constitutive tonic signaling activity.

Flow cytometry analysis revealed variable levels but ubiquitous surface expression of B7-H3 on various DIPG cell lines including DIPG-C1, DIPG2, DIPG4, DIPG007, DIPG13, DIPG21, and DIPG36 (Figure S1A, B). This indicates that DIPG cells express B7-H3, making them suitable targets for B7-H3-directed CAR-T therapy.

Next, we assessed the cytotoxicity of these CAR-T cells on DIPG cells in culture. B7H3.BC CAR-T cells more efficiently killed DIPG36 cells carrying a luciferase reporter than MGA271 or Hu8H9 CAR-T cells. After 48 hours of co-culture at effector (T cells)-to-target (tumor cells) (E:T) ratios of 1:2 and 1:4, B7H3.BC CAR-T cells most effectively cleared tumor cells (Figure 1C), indicating superior killing efficacy. Consistent with these findings, B7H3.BC CAR-T cells caused more pronounced reductions in DIPG36, DIPG13, DIPG-C1, and DIPG21 cell numbers than MGA271 and Hu8H9 CAR-T cells over a 5-day co-culture period (Figure 1D, E and Figure S1C). Cytokine release assays showed that B7H3.BC CAR-T cells secreted significantly higher levels of IFNγ and IL-2 than MGA271 and Hu8H9 CAR-T cells after 24 hours of co-culture with multiple DIPG cells (DIPG36, DIPG13, and DIPG21) (Figure 1F). This suggests that the enhanced cytotoxicity observed with B7H3.BC CAR-T cells is associated with increased T functions in the presence of tumor cells.

Using fluorophore AF647–conjugated B7-H3 protein as a substrate, we found that B7H3.BC CAR-T cells bound B7-H3 with comparable capacity to MGA271 and Hu8H9 CAR-T cells (Figure S1D). Despite this equivalent antigen-binding capacity, B7H3.BC CAR-T cells with subdued tonic signaling exhibit superior killing efficacy against DIPG cells compared with MGA271 and Hu8H9 CAR-T cells.

### Tonic signaling restraint sustains stem-like phenotypes and limits differentiation

To investigate the differentiation states of B7H3 CAR-T cells, we performed flow cytometry analysis on day 14 of culture. Surface expression of CD62L and CD45RO was used to distinguish naïve T cells/stem cell memory T cells (T_N_/T_SCM_, CD62L⁺CD45RO⁻), central memory T cells (T_CM_, CD62L⁺CD45RO⁺), effector memory T cells (T_EM_, CD62L⁻CD45RO⁺), and effector T cells (T_EFF_, CD62L⁻CD45RO⁻) ^28,29^. Flow cytometry assays indicate distinct differentiation profiles across the different CAR-T constructs (Figure S2A).

Analysis of T cell subpopulations revealed that cultures of B7H3.BC CAR-T cells had a significantly higher proportion of stem-like naïve/stem cell memory T cells than MGA271 and Hu8H9 CAR-T cell cultures (Figure S2B, C). Specifically, B7H3.BC CAR-T cells were enriched with stem-like cells, whereas MGA271 and Hu8H9 CAR-T cells were enriched with the effector memory phenotype, indicative of a more differentiated state (Figure S2C).

Assessment of stem-like and memory markers (T_N_/T_SCM_) further supported these findings. Flow cytometry analysis of CD62L, CCR7, and CD45RA, which are stem-like markers, and CD45RO, a T cell memory marker ^28,29^ revealed that B7H3.BC CAR-T cells had higher expression of CD62L, CCR7, and CD45RA and lower expression of CD45RO compared to MGA271 and Hu8H9 CAR-T cells on day 14 of culture (Figure S2D, E). There were minimal differences in levels of these markers between the CAR-T cell groups at the early timepoint day 7 (Figure S2F, G), suggesting that the differentiation of MGA271 and Hu8H9 CAR-T cells becomes more apparent after this time point. Together, these data indicate that B7H3.BC CAR-T cells maintain a stem-like, less differentiated phenotype, which may contribute to their persistence and therapeutic efficacy.

### Restrained tonic signaling limits exhaustion in CAR-T cells

To evaluate the impact of tonic signaling on the CAR-T cell exhaustion, we analyzed the expression of T cell exhaustion and activation markers after 14 days of culture. Flow cytometry analysis showed that B7H3.BC CAR-T cells had lower levels of surface expression of exhaustion markers (PD-1, TIM3, and LAG3) ^30^ and T cell activation markers (CD25 and CD69) than did MGA271 and Hu8H9 CAR-T cells (Figure 2A, B). Compared to MGA271 and Hu8H9 CAR-T cells, B7H3.BC CAR-T cells exhibited a reduced exhaustion phenotype, as indicated by a significantly lower percentage of TIM3⁺LAG3⁺ double-positive cells (Figure 2C). In addition, B7H3.BC CAR-T cell cultures displayed fewer apoptotic cells (Figure 2D) and greater proliferative capacity (Figure 2E). This suggests that B7H3.BC CAR-T cells exhibit enhanced functional and proliferative profiles with reduced exhaustion and apoptosis relative to MGA271 and Hu8H9 CAR-T cells.

**Figure 2.**
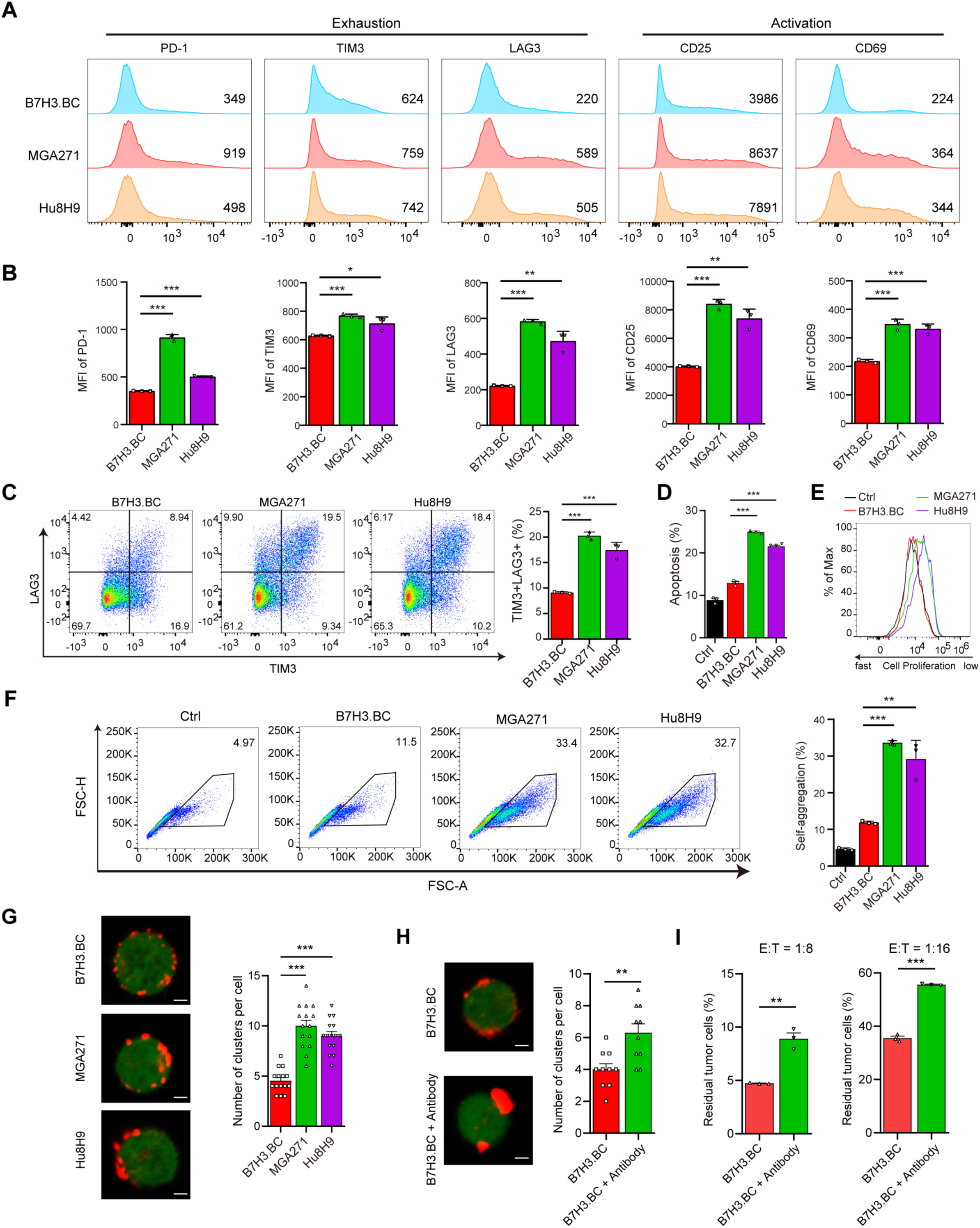
B7H3.BC CAR-T cells exhibit attenuated antigen-independent T cell exhaustion. **(A)** Flow cytometric analysis of exhaustion markers (PD-1, TIM3, and LAG3) and activation markers (CD25 and CD69) on B7-H3 CAR-T cells after 14 days of culture. Representative of three donors. **(B)** MFIs of indicated markers on B7-H3 CAR-T cells after 14 days of culture. **(C)** Representative flow cytometry plots (left) and quantification (right) of exhaustion of B7-H3 CAR-T cells based on TIM3 and LAG3 expression. Representative of three donors. **(D)** Quantification of apoptosis in B7-H3 CAR-T cell cultures on day 14. Representative of three donors. **(E)** Representative analysis of ViaFluor 405 staining as a measure of cell proliferation on day 14 in B7-H3 CAR-T cells and control T cell cultures. Representative of three donors. **(F)** Representative flow cytometry analysis (left) and quantification of aggregated clusters per cells (right) on day 7 in CAR-T and control T cell cultures. Representative of three donors. **(G)** (Left) Representative confocal images showing the surface distribution of CAR molecules on indicated B7-H3 CAR-T cells. Scale bars = 5 µm. (Right) Quantification of CAR clusters per cell, based on analysis of 15 cells per group. **(H)** (Left) Representative confocal images showing the surface distribution of CAR molecules on untreated or antibody-treated B7H3.BC CAR-T cells. Scale bars = 5 µm. (Right) Quantification of CAR clusters per cell, based on analysis of 10 cells per group. **(I)** Percent survival of DIPG36 cells incubated with untreated or antibody-treated B7H3.BC CAR-T cells relative to sample incubated with control T cells assessed by luciferase assays after 2 days of co-culture. CAR-T effector to tumor cell ratios of 1:8 and 1:16. Data are means ± SEM from triplicate wells. Representative of three donors. For panels **B-D** and **F-I**, unpaired two-tailed Student’s *t*-test. *P < 0.05, **P < 0.01, ***P < 0.001.

To investigate the basis of these functional differences, we next evaluated CAR clustering on cell membranes as a functional readout of tonic signaling ^31^. After 7 days of culture, MGA271 and Hu8H9 CAR-T cells had significantly higher levels of spontaneous CAR-T aggregation than did B7H3.BC CAR-T cells or control T cells, which were transduced with empty vector, as measured by flow cytometry (Figure 2F). Although the B7-H3 CARs exhibited comparable expression levels (Figure S3A), MGA271 and Hu8H9 CAR-T cells exhibited pronounced clustering, whereas B7H3.BC CAR-T cells had a more even CAR distribution (Figure 2G). This is consistent with the subdued tonic signaling observed in B7H3.BC CAR-T cells, in contrast to the elevated tonic signaling seen in MGA271 and Hu8H9 CAR-T cells.

To assess the impact of CAR clustering on CAR-T function, we treated B7H3.BC CAR-T cells with an anti-G4S antibody to induce clustering on the T cell membrane. Antibody-treated cells formed more prominent CAR clusters than untreated controls (Figure 2H). When evaluating their cytotoxicity against luciferase-expressing DIPG36 cells, we found that the antibody-induced CAR clustering reduced the killing capacity of B7H3.BC CAR-T cells (Figure 2I), indicating that increased CAR clustering impairs CAR-T function.

To further explore the structural basis underlying CAR clustering, we used AlphaFold2-Multimer to predict scFv dimer structures (Figure S3B) and assess CAR dimerization propensity ^32-34^. Rosetta scores indicated weaker dimerization for B7H3.BC compared to MGA271 and Hu8H9 (Figure S3C). Consistently, steered molecular dynamics simulations ^35,36^ showed greater pulling energy, and thus stronger dimerization, for MGA271 and Hu8H9 (Figure S3D, E). Thus, compared to MGA271 and Hu8H9, these results indicate that the B7H3.BC has a lower clustering propensity, which may contribute to reduced constitutive T cell activation and exhaustion.

### Tonic-subdued CAR-T cells maintain long-term efficacy after tumor rechallenge

To evaluate the long-term proliferative capacity and antitumor activity of B7-H3-targeting CAR-T cells, we performed three-round co-culture rechallenge assays ^37^ using DIPG13 cells and CAR-T cells at an effector-to-target ratio of 1:4 (Figure 3A). Flow cytometry was used to quantify the proportion of mCherry-labeled DIPG13 cells (tumor) and GFP-labeled CAR-T cells at the end of each co-culture round. After the first round of co-culture, all three CAR-T cells reduced the population of DIPG13 cells compared to control T cells (Figure 3B). In the second co-culture round, B7H3.BC CAR-T cells showed higher numbers of GFP⁺ CAR-T cells and fewer mCherry⁺ tumor cells compared to MGA271 and Hu8H9 CAR-T cultures (Figure 3B, C; Figure S4A). This trend persisted through the third round of co-culture, during which B7H3.BC CAR-T cells continued to expand more robustly and to eliminate tumor cells more effectively than MGA271 or Hu8H9 CAR-T cells (Figure 3B, C and Figure S4B, C). The enhanced expansion of B7H3.BC CAR-T cells after multiple rounds of tumor exposure indicates superior persistence and resistance to exhaustion.

**Figure 3.**
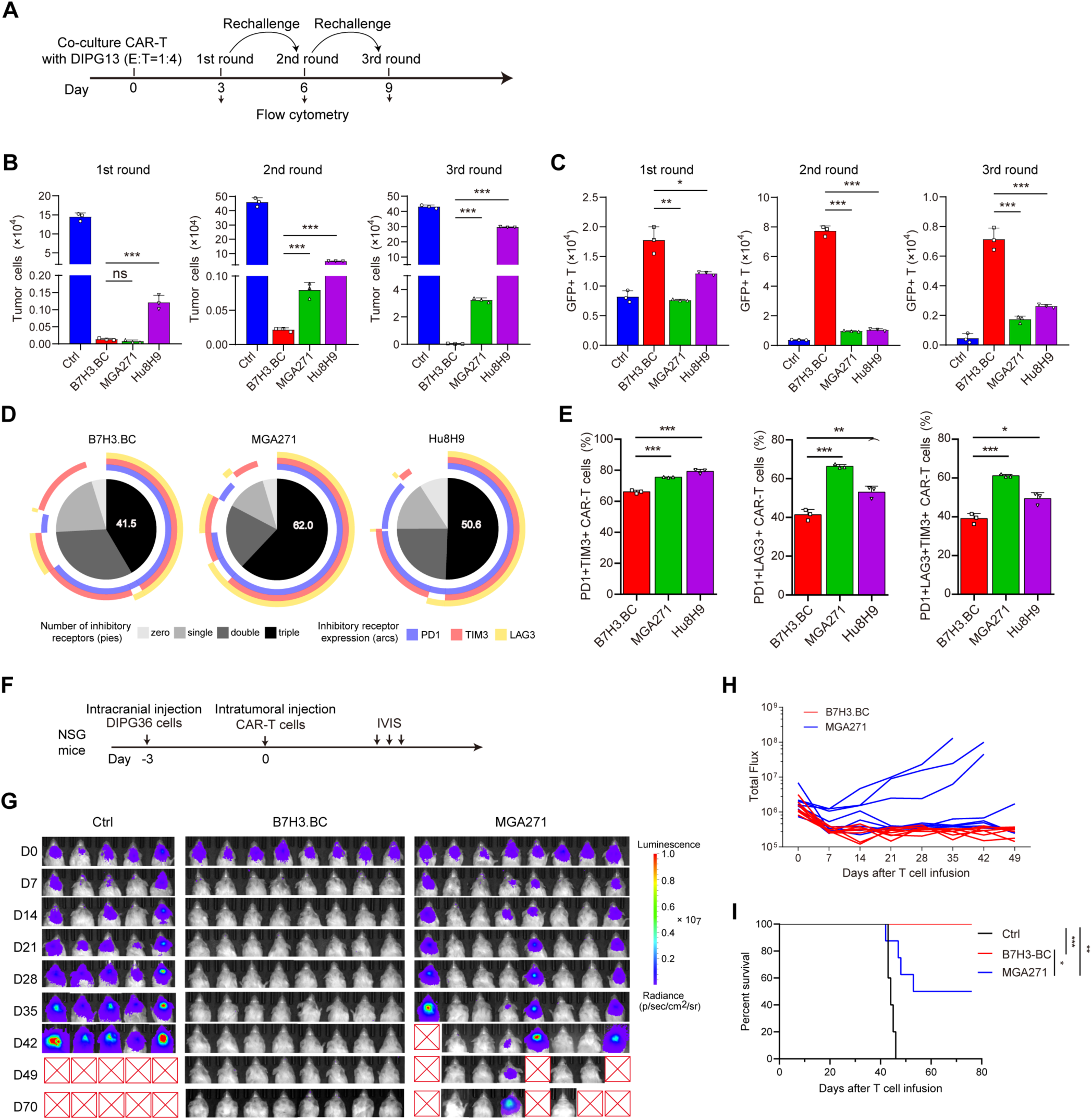
B7H3.BC CAR-T cells exhibit long-term antitumor activity in vitro and in vivo. **(A)** Schematic overview of the three-round co-culture experiment using DIPG13 cells and CAR-T cells. **(B, C)** Quantification of mCherry-labeled DIPG13 cells (C) and GFP-labeled CAR-T cells (D) after indicated rounds of co-culture. CAR-T cells and tumor cells were co-cultured at a ratio of 1:4. Representative of three donors. **(D)** SPICE analysis of the expression of exhaustion markers PD-1, TIM3, and LAG3 in CAR-T cells after the second round of co-culture. **(E)** Percentages of exhausted populations after the second round of co-culture. **(F-I)** A total of B7H3.BC CAR-T cells, MGA271 CAR-T cells, or control T cells were infused intratumorally into NSG mice 3 days after intracranial injection of 1 × 10⁵ DIPG36 cells. IVIS was used to quantify bioluminescence over time. **F)** Schematic overview of the experiment. **G)** Bioluminescence images showing tumor growth (Ctr, n=5 mice; B7H3.BC, n=8 mice; MGA271, n=8 mice). **H)** Bioluminescence flux per individual mouse over time. **I**) Kaplan-Meier survival curve of NSG mice. For panels **B**, **C** and **E**, unpaired two-tailed Student’s *t*-test. *P < 0.05, **P < 0.01, ***P < 0.001; ns, not significant. In panel **I**, the log-rank test. **P < 0.01, ***P < 0.001.

To investigate the exhaustion status of CAR-T cells following repeated antigen exposure, we examined the expression of exhaustion markers PD-1, TIM3, and LAG3 ^38,39^ after the second round of co-culture using flow cytometry. Levels of co-expression of these exhaustion markers were lower on B7H3.BC CAR-T cells than on MGA271 and Hu8H9 CAR-T cells. Quantification of the different exhausted subpopulations confirmed that B7H3.BC CAR-T cells had lower proportions of PD-1⁺TIM3⁺LAG3⁺ triple-positive cells, indicative of reduced exhaustion, than did the other B7-H3-CAR-T cell lines (Figure 3D, E). These findings demonstrate that B7H3.BC CAR-T cells possess enhanced proliferative capacity and long-term antitumor activity in repeated rechallenge assays, which correlates with a reduced exhaustion phenotype and superior persistence compared to MGA271 and Hu8H9 CAR-T cells.

### CAR-T cells with subdued tonic signaling enhance infiltration and cytotoxicity in DIPG spheroids

To evaluate the cytotoxic activity of B7-H3-targeted CAR-T cells in a context that mimics the three-dimensional (3D) architecture of tumors, we co-cultured CAR-T cells with patient-derived DIPG spheroids. Spheroids were seeded in 8-well glass chambers and allowed to establish for 3 days prior to the addition of CAR-T cells (Figure S5A). B7H3.BC CAR-T cells caused a progressive and substantial reduction in organoid size and fluorescence intensity over 5 days, indicative of tumor cell clearance (Figure S5B).

To investigate dynamics of CAR-T cells during disruption of solid tumor architecture, we co-cultured DIPG spheroids with either B7H3.BC or MGA271 CAR-T cells. While both disrupted the spheroid structure, B7H3.BC CAR-T cells more effectively infiltrated the spheroids and showed greater tumor cell targeting (Figure S5C, D). These findings suggest that B7H3.BC CAR-T cells possess enhanced cytotoxic activity compared to MGA271 CAR-T cells against DIPG-derived tumor spheroids.

### Potent antitumor efficacy by locoregional delivery of tonic-subdued CAR-T cells in DIPG xenografts

To evaluate the *in vivo* antitumor efficacy of B7H3.BC CAR-T cells, we employed an orthotopic xenograft model using luciferase-expressing DIPG36 cells in immunodeficient NSG mice. Mice were infused intracranially with CAR-T cells or control T cells after tumor cell injection (Figure S6A). Bioluminescence imaging revealed marked suppression of tumor progression in mice treated with B7H3.BC CAR-T cells compared to mice treated with control T cells, which had been transduced with empty vector (Figure S6B). Tumor burden remained minimal in CAR-T–treated animals over the course of the study, while control mice exhibited rapid tumor growth. Kaplan-Meier analysis showed a significant extension of survival in the B7H3.BC CAR-T cell-treated group relative to controls (Figure S6C). These findings suggest that B7H3.BC CAR-T cells exert potent antitumor activity in the xenograft model.

To further compare the *in vivo* efficacies of B7H3.BC and MGA271 CAR-T cells, we utilized the DIPG36 xenograft model. DIPG36 cells were injected three days prior to intratumoral injection of CAR-T cells or control T cells (Figure 3F). We observed markedly greater tumor reduction in mice treated with B7H3.BC CAR-T cells than in those that received MGA271 CAR-T cells (Figure 3G). Tumor burden was significantly lower over time in the group of mice treated with B7H3.BC CAR-T cells compared to the MGA271 CAR-T cell-treated group (Figure 3H). Treatment with B7H3.BC CAR-T cells significantly prolonged survival compared to the MGA271 CAR-T cell group (Figure 3I), indicating potent antitumor activity in the xenograft models.

### mRNA-based tonic-subdued CAR-T cells exhibit potent antitumor activity against DIPG

mRNA-based CAR-T cell therapy offers several advantages over traditional viral vector-based approaches, primarily due to its transient and negligible genomic integration, leading to improved safety and flexibility ^40-42^. To evaluate the antitumor efficacy of B7H3.BC CAR-T cells engineered using the mRNA-based strategy, we first transfected B7H3.BC stable mRNA-CAR into T cells using the MaxCyte Flow-Electroporation method and assessed their killing ability against DIPG cells in co-culture experiments. The B7H3.BC mRNA-based CAR-T cells efficiently eliminated DIPG13, DIPG21, and DIPG36 cells over a 5-day co-culture period in 1:1 culture, whereas control GFP-mRNA expressing T cells, showed minimal tumor cell clearance under these conditions (Figure S7A, B). The progressive reduction in mCherry-expressing DIPG cells and expansion of GFP-expressing CAR-T cells over time highlighted the potent cytotoxic activity of B7H3.BC mRNA-based CAR-T cells.

To assess the antitumor activity of B7H3.BC mRNA-based CAR-T cells *in vivo*, we used an orthotopic DIPG36 xenograft model in NSG mice. Three days after intracranial injection of DIPG36 cells, mice were infused intravenously with 1 × 10⁷ B7H3.BC mRNA-based CAR-T cells or control T cells (Figure S7C). Bioluminescence imaging revealed significant tumor reduction in mice treated with B7H3.BC mRNA-based CAR-T cells compared to mice treated with control T cells (Figure S7D). Quantification of tumor burden over time showed that in mice treated with B7H3.BC mRNA-based CAR-T cells there was significantly slower tumor growth compared to control mice (Figure S7E). This reduction in tumor burden correlated with prolonged survival. B7H3.BC mRNA-based CAR-T-treated group showed a significant survival benefit compared to the control group (Figure S7F). Histological analysis of treated brains revealed tumor clearance in mice receiving B7H3.BC mRNA-based CAR-T cells, in contrast to the high tumor burden observed in control T cell-treated mice (Figure S7G). These observations indicate that B7H3.BC mRNA-based CAR-T cells exert potent and sustained antitumor activity against DIPG both *in vitro* and *in vivo*, resulting in tumor reduction and improved survival outcomes.

### Subdued tonic signaling imprints transcriptional and epigenomic programs supportive of CAR-T functionality

To investigate potential mechanism underlying the enhanced antitumor efficacy of B7H3.BC CAR-T cells, we performed RNA sequencing and epigenomic profiling of B7H3.BC, MGA271, and Hu8H9 CAR-T cells and control T cells transduced with empty vector as well as untransduced T cells. Principal component analysis of RNA-seq data revealed distinct clustering of B7H3.BC CAR-T cells from other cells (Figure S8A), indicative of unique transcriptional programs in B7H3.BC CAR-T cells. Differential gene expression analysis showed that the gene expression signature of B7H3.BC CAR-T cells was distinct from MGA271 and from Hu8H9 CAR-T cells (Figure S8B). Notably, Gene Ontology analysis revealed significant downregulation of pathways associated with TCR receptor tonic signaling, T cell activation, cell-cell adhesion and upregulation of pathways associated with chemotaxis, proliferation, and cell survival in B7H3.BC CAR-T cells relative to MGA271 and Hu8H9 CAR-T cell counterparts (Figure S8C). Moreover, analysis of transcription regulatory programs revealed that the downregulated genes are the targets of tonic signaling-associated transcription factors such as *NFKB1, RELA,* and *STAT3* (Figure S8D).

Genes related to tonic signaling-associated signaling pathways including the MAPK/ERK (e.g., *FGF2*, *NOD2*), PI3K (e.g., *RAMP3*, *SRC*), TNF (e.g., *TRAF1*, *TNFRSF18*), JNK (e.g., *MAP3K6*, *RIPK2*), and STAT3 (e.g. *IL23R*, *IL6R*) pathways and to apoptosis-associated pathways (e.g., *DAPK1*, *RPS6KA2*) were expressed at lower levels in B7H3.BC CAR-T cells than in MGA271 and Hu8H9 CAR-T cells (Figure S8E). Pathway enrichment analysis also indicated attenuation of tonic signaling and T cell exhaustion-related pathways among the downregulated genes in the B7H3.BC CAR-T group (Figure S8F). Consistently, we observed reduced enrichment of the TNF/NF-κB signaling hallmark signature (Figure S8G), which is associated with chronic inflammation and T cell dysfunction ^43,44^.

Transcriptome analysis of T cell functional states indicated that B7H3.BC CAR-T cells also exhibited reduced expression of exhaustion- and effector-associated genes and elevated expression of stemness markers such as *KLF2*, *SELL* (which encodes CD62L), and *MYC* compared to MGA271 and Hu8H9 CAR-T cells (Figure S8H). GSEA further revealed enriched KLF2 stemness targets and reduced exhaustion signatures in B7H3.BC CAR-T cells (Figure S8I). These findings are consistent with the enhanced stem-like, less differentiated, and less exhausted phenotype observed in B7H3.BC CAR-T cells.

T cell metabolic profiles are closely tied to their functional states, with oxidative phosphorylation (OXPHOS) linked to stem-like properties and glycolysis associated with more differentiated phenotypes ^45-47^. GSEA and expression profiling showed that OXPHOS pathway genes (e.g., *NDUFB2, NDUFS6, COX7B*) were upregulated in B7H3.BC CAR-T cells compared to MGA271 and Hu8H9 CAR-T cells (Figure S8J, K), consistent with their stem-like phenotype. Conversely, B7H3.BC CAR-T cells exhibited reduced enrichment of glycolysis-related genes (e.g., *ALDOC, PKM, PFKM*), reflecting a less differentiated state. Additionally, genes involved in the cGAS–STING signaling pathway, an innate immune sensing cascade, were elevated in B7H3.BC CAR-T cells (Figure S8L-N). These findings suggest that B7H3.BC CAR-T cells possess an enhanced stem-like transcriptional program characterized by restrained activation, reduced tonic signaling, and increased innate immune sensing.

To further investigate the epigenetic enhancer landscape, we performed CUT&Tag profiling of H3K27ac, a marker of active regulatory enhancer regions ^48^, in control T cells and B7-H3 CAR-T cells. Consistent with the RNA-seq data, the downregulated genes in B7H3.BC CAR-T exhibited significantly lower H3K27ac enrichment near the transcription start site (TSS) compared to MGA271 and Hu8H9 CAR-T cells (Figure S9A). B7H3.BC CAR-T cells showed reduced H3K27ac signal intensity at loci associated with T cell exhaustion (e.g., *BATF3*, *TOX2*) and tonic signaling (e.g., *DAPK1*, *IL23R*) (Figure S9B, C), while H3K27ac signals at stemness-associated loci (e.g., *KLF2*, *MYC*) were elevated (Figure S9D). In contrast, the H3K27ac signal levels at housekeeping genes (e.g., *GAPDH*, *TUBA1A, TBP, ACTB*) remained comparable across the CAR-T cell types (Figure S9E). Together, these analyses indicate that B7H3.BC CAR-T cells adopt a transcriptional and epigenetic state of heightened stemness, chemotaxis, and innate immune sensing, with reduced tonic signaling and exhaustion.

### Single-cell transcriptomics reveal tonic signaling drives early progenitor-exhausted-like states in CAR-T Cells

To identify transcriptional signatures linked to CAR-T function, we performed clustering analysis of differentially expressed genes across untransduced/control T cells (CTR), B7H3.BC CAR-T cells, and other CAR-T constructs (MGA271 and Hu8H9) (Figure 4A; Figure S10A). Pathway enrichment revealed discrete gene expression modules including a tonic signaling–associated signature (CAR-Ton), a CAR-T effector-memory signature (CAR-Tem), and a CAR-T inflammatory signature (CAR-Infl) (Figure 4B and Figure S10B; Supplementary Table 1). Single-sample GSEA (ssGSEA) analysis further showed that B7H3.BC CAR-T cells exhibit markedly lower CAR-Ton scores but higher CAR-Tem and CAR-Infl scores than MGA271 and Hu8H9 CAR-T cells (Figure 4C; Figure S10C).

**Figure 4.**
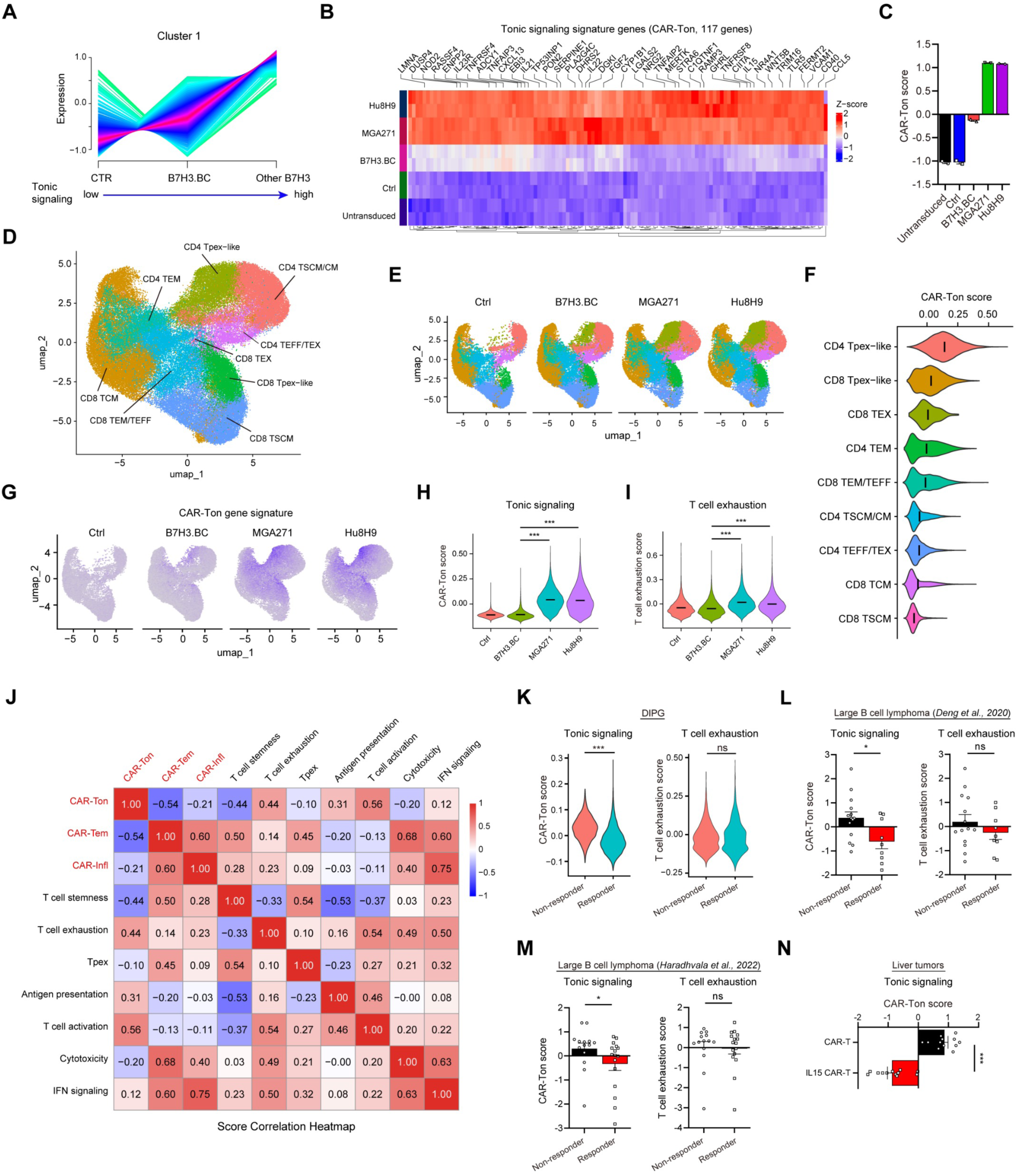
Tonic signaling score as a predictive biomarker of CAR-T efficacy. **(A)** Clustering plot showing scaled expression changes of genes in CTR (untransduced and Ctrl), B7H3.BC CAR-T, and Other B7-H3 CAR-T (MGA271 and Hu8H9). Gene expression values were normalized to z-scores for visualization. **(B)** Heatmaps depicting differentially expressed genes associated with the tonic signaling score. The representative genes were showed at the top. **(C)** The tonic signaling (CAR-Ton) score in untransduced T cells, control T cells, and B7H3.BC, MGA271 and Hu8H9 CAR-T cells. The scores were calculated by the ssGSEA method based on the corresponding gene list. **(D)** UMAP visualization of scRNA-seq data from 87,028 cells, including control T cells (Ctrl), and B7H3.BC, MGA271, and Hu8H9 CAR-T cells on day 12 of culture. Nine clusters are indicated by different colors. **(E)** Side-by-side UMAP visualization showing the distribution of nine clusters in Ctrl (15,928 cells), B7H3.BC (25,666 cells), MGA271 (23,436 cells) and Hu8H9 (21,998 cells) CAR-T cells. **(F)** Violin plot of CAR-Ton scores in different clusters. The crossbar represents mean value of the scores. The clusters were arranged based on the mean values of the scores from high to low. **(G)** Feature plot of tonic signaling (CAR-Ton) score in control T and CAR-T cells. **(H)** Violin plot of CAR-Ton scores in control and B7-H3 CAR-T cells. The crossbar represents mean value of the scores. The scores were calculated by the AddModuleScore function of Seurat. **(I)** Violin plot of T cell exhaustion scores in control and B7-H3 CAR-T cells. The crossbar represents mean value of the scores. The scores were calculated by the AddModuleScore function of Seurat. **(J)** The score correlation heatmap using the data from Dataset (*Deng et al., 2020*). The scores were calculated by the ssGSEA method based on the corresponding gene lists. **(K)** Comparison of the CAR-Ton score (left) or T cell exhaustion score (right) in the GD2 CAR-T non-responder and responder using the data from Dataset (*Majzner et al., 2022*) in DIPG. The scores were calculated by the AddModuleScore function of Seurat. **(L)** Comparison of the CAR-Ton scores or T cell exhaustion scores in CAR-T non-responders and responders in using the data from Dataset (*Deng et al., 2020*; non-responder, n = 14; responder, n = 9) in large B cell lymphoma. The scores were calculated by the ssGSEA method. **(M)** Comparison of the CAR-Ton scores or T cell exhaustion scores in CAR-T non-responders and responders in using the data from Dataset (*Haradhvala et al., 2022*; non-responder, n = 14; responder, n = 16) in large B cell lymphoma. The scores were calculated by the ssGSEA method. **(N)** Comparison of the CAR-Ton scores in CAR-T vs IL-15 CAR-T patients using the data from Dataset (*Steffin et al., 2024*; CAR-T, n = 12; IL-15 CAR-T, n = 12) in liver tumors. The scores were calculated by the ssGSEA. For panel **H** and **I**, Kruskal-Wallis test followed by pairwise Wilcoxon rank-sum tests (BH-adjusted). For panel **K**, Wilcoxon rank-sum test. For panels **L**-**N**, unpaired two-tailed Mann–Whitney test. *P < 0.05, **P < 0.01, ***P < 0.001; ns, not significant.

To assess the impact of tonic signaling on CAR-T cell populations, we conducted single-cell transcriptomics profiling on sorted GFP⁺ control and CAR-T cells. UMAP projection identified nine transcriptionally distinct cell populations (Figure 4D) defined by cluster-specific marker genes (Figure S11A). Strikingly, progenitor-exhausted–like (Tpex-like) cells among CD4⁺ and CD8⁺ populations (*TCF7*^high^ *CCR7*^high^ *TOX2*^high^ *KLF2*^low^) were markedly enriched in MGA271 and Hu8H9 CAR-T cells compared with B7H3.BC CAR-T cells (Figure 4E; Figure S11B), suggesting that high tonic-signaling MGA271 and Hu8H9 CAR-T cells are more prone to entering a progenitor-exhaustion state than B7H3.BC CAR-T cells. These Tpex-like cells also exhibited the highest CAR-Ton signature scores among the T cell populations (Figure 4F). Flow cytometry confirmed the increased frequencies of the cells expressing Tpex markers TCF1/TIM3^49^ (Figure S11C). Furthermore, compared to B7H3.BC CAR-T cells, MGA271 and Hu8H9 cells showed elevated CAR-Ton and T cell exhaustion scores (Figure 4G–I) alongside reduced CAR-Tem scores (Figure S11D). Collectively, these findings indicate that elevated tonic signaling in CAR-T cells results in differentiation toward a progenitor-exhausted–like state, thereby potentially leading to T cell exhaustion and constraining their long-term persistence.

### A tonic signaling signature score predicts clinical outcome and CAR-T cell efficacy

To assess the clinical relevance of the CAR-T tonic signature (CAR-Ton) score, we analyzed publicly available scRNA-seq datasets from CAR-T clinical studies, including DIPG, large B cell lymphomas, and liver cancer ^9,50,51^. We evaluated clinical outcomes in responders and non-responders by comparing the CAR-Ton score with previously defined T-cell functional gene signatures, including T-cell exhaustion ^52,53^, stemness ^54,55^, activation ^56,57^, antigen presentation ^58,59^, cytotoxicity ^60,61^, IFN signaling ^62,63^, and Tpex states ^49^ (Supplementary Table 2). Correlation analysis showed that the CAR-Ton score positively correlated with T-cell activation, exhaustion, and antigen presentation scores, but negatively correlated with CAR-Tem and T-cell stemness scores (Figure 4J), suggesting that the tonic signaling score represents an integrated signature of these T-cell functional properties.

CAR-Ton scores were significantly higher in non-responders compared to responders in the GD2 CAR-T clinical trials for DIPG/DMG^9^ (Figure 4K). Consistently, Elevated CAR-Ton scores were consistently observed in non-responders across two independent CD19 CAR-T clinical studies in large B cell lymphoma ^50,51^ (Figure 4L, M). In contrast, T cell exhaustion and stemness scores, commonly used metric for assessing T cell function, showed no significant difference between responders and non-responders in these clinical studies (Figure 4K-M and Figure S12A). These suggest that traditional T cell exhaustion or stemness signature is insufficient to predict CAR-T clinical responses. Likewise, Tpex, CAR-Tem, and pro-inflammatory CAR-Infl scores also showed no significant differences between responders and non-responders (Figure S12B-D). Receiver operating characteristic (ROC) analysis further confirmed the predictive value of the CAR-Ton score, with AUC values consistently exceeding 0.7 across two independent trial datasets in large B cell lymphoma (Figure S12E).

To further link tonic signaling score with antitumor activity, we analyzed the single cell RNA-seq dataset from CAR-T clinical trials in liver tumors ^64^, in which IL-15–engineered CAR-T cells displayed enhanced expansion, persistence, and tumor control. CAR-Ton scores were markedly lower in IL-15 CAR-T cells compared to conventional CAR-T cells (Figure 4N). Additionally, tonic signaling–associated genes were consistently downregulated in IL-15 CAR-T cells (Figure S12F). Collectively, these findings support a model in which restrained tonic signaling prevents CAR-T cells from differentiating into progenitor-exhausted dysfunctional states, positioning the CAR-Ton signature as a robust correlate of both clinical outcome and antitumor activity.

## DISCUSSION

### CAR-T tonic signaling as a mechanistic determinant of anti-tumor efficacy in DIPG/DMG

DIPG/DMG remains one of the most lethal pediatric brain tumors, and despite the promise of CAR-T cell therapy, clinical responses are inconsistent, often limited by premature exhaustion and poor persistence. Our study identifies tonic signaling as a central determinant of CAR-T efficacy in DIPG/DMG. By developing and systematically interrogating multiple B7-H3–directed CAR constructs under clinical investigation, we show that subdued tonic signaling confers markedly superior antitumor activity, persistence, and resistance to exhaustion in *in vitro* and *in vivo* DIPG models, despite targeting the same antigen with comparable binding capacity.

Although antigen-independent tonic signaling is recognized as influencing CAR-T function, its quantitative impact on T-cell fate and function as well as its mechanistic links to the transcriptional and epigenetic programs governing CAR-T durability in DMG/DIPG remain undefined. Our systematic comparative analyses of clinically relevant B7-H3 CARs reveal that attenuated tonic signaling enhances persistence, infiltration, and therapeutic efficacy of B7-H3 CAR-T cells in patient-derived DIPG/DMG models. We further delineate a transcriptional and epigenetic landscape that mechanistically links tonic signaling to progenitor-exhausted–like states and identify a CAR-T tonic signaling–associated signature as a robust predictor of CAR-T efficacy, outperforming conventional exhaustion or stemness markers across multiple independent clinical trials, including those in DIPG and other tumors. Thus, these findings advance our understanding beyond the general recognition of tonic signaling to a mechanistically defined and clinically predictive framework for informing potential therapeutic outcomes.

### Identification of tonic-subdued B7-H3 CAR with enhancing antitumor efficacy across tumor models

The enhanced antitumor activity of B7H3.BC CAR-T cells with restrained tonic signaling was consistently observed across diverse experimental models. In *in vitro* tumor cell killing assays and tumor rechallenge experiments, B7H3.BC CAR-T cells showed superior cytotoxicity against multiple DIPG cell lines, coupled with more sustained proliferation and greater resistance to functional exhaustion compared to the other CAR-T constructs tested. In 3D tumor spheroid models, they exhibited markedly improved infiltration and clearance of tumor cells, reflecting enhanced migratory capacity and persistence within a complex tumor architecture. Importantly, these in vitro advantages translated into potent therapeutic activity in vivo, where B7H3.BC CAR-T cells achieved significant tumor regression and prolonged survival in patient-derived orthotopic DIPG xenograft models. Together, these results indicate that B7H3.BC CAR-T cells with subdued tonic signaling are functionally enhanced and capable of overcoming key barriers of persistence, exhaustion, and tumor infiltration that limit the efficacy of other CAR-T cell designs in DIPG.

We found that B7H3.BC CAR-T cells exhibit markedly attenuated antigen-independent tonic signaling as evidenced by reduced basal signaling activity across the signaling effectors (e.g., pAKT, pS6, pSTAT3, pERK). This subdued tonic signaling was correlated with a less differentiated, stem-like phenotype, and diminished expression of exhaustion markers (PD-1, TIM3, LAG3), the hallmarks of early T cell dysfunction ^65,66^. Moreover, B7H3.BC CAR-T cells have less tonic signaling-associated CAR clustering on the membrane than did the other CAR-T cells evaluated. Structural modeling and molecular dynamics simulations further implicated reduced CAR membrane clustering as a distinguishing feature of B7H3.BC CARs. Notably, experimentally induced clustering impaired their efficacy, suggesting that limiting CAR receptor self-association is critical for maintaining functional potency.

The utility of B7H3.BC CAR-T cells was extended to an mRNA-based delivery platform, which offers advantages for transient CAR expression with reduced risk of insertional mutagenesis and greater control over therapeutic windows ^42,67,68^. B7H3.BC CAR-T cells maintained superior antitumor activity *in vitro* and *in vivo* when expressed transiently via mRNA electroporation, supporting the feasibility of non-integrating CAR strategies in DIPG/DMG. mRNA-based CAR-T cells may permit more flexibility and safer delivery platforms, especially in vulnerable patient populations such as children. Transient CAR expression via mRNA delivery allows for iterative dosing and rapid discontinuation in the event of adverse effects ^42,67,68^, a potential beneficial feature especially relevant in the sensitive context of CNS malignancies.

### Multi-omics profiling reveals transcriptomic and epigenetic programs underling CAR functionality

Our integrated transcriptomic and epigenomic profiling revealed that reduced tonic signaling reprograms CAR-T cells toward a transcriptional and epigenetic state with enhanced functionality. B7H3.BC CARs downregulated exhaustion-associated gene programs while upregulating stemness, OXPHOS metabolism, and innate immune sensing. Epigenomic enhancer profiling confirmed an epigenetic landscape permissive to memory and self-renewal. Thus, reduced tonic signaling in B7H3.BC CAR-T cells not only limits exhaustion but may also preserve an epigenetic landscape permissive to T cell self-renewal programs.

Our single-cell transcriptomic analyses further linked high tonic signaling to progenitor-exhausted differentiation trajectories. Elevated tonic signaling, as observed in MGA271 and Hu8H9 constructs, skews T-cell populations toward a progenitor-exhausted-like state characterized by heightened expression of exhaustion markers and diminished stemness signatures. This differentiation trajectory is associated with reduced persistence potential, contributing to the inferior therapeutic performance of high tonic-signaling CAR-T cells. In contrast, low-tonic B7H3.BC CAR-T cells retain greater stemness and exhibit reduced exhaustion. Thus, our integrated single-cell and multi-omic profiling identify antigen-independent tonic signaling as a key driver of transcriptional and epigenetic programs that govern CAR-T durability and efficacy.

### A CAR-T tonic signaling framework for predicting therapeutic outcomes

Our integrated multi-omic analysis defines a tonic signaling–associated gene signature that is predictive of clinical responses. Across multiple independent clinical trials, including DIPG and other cancer types ^9,50,51^, antigen-independent tonic signaling signature scores robustly predicted response to CAR-T therapy, outperforming traditional downstream markers such as T cell exhaustion signatures ^14,15^. Although the precise mechanisms remain to be defined, tonic signaling may function as an upstream intrinsic driver that coordinates activation, differentiation, and survival programs, thereby imprinting transcriptional and epigenetic states that govern CAR-T persistence, exhaustion, and efficacy. By contrast, exhaustion and stemness markers largely reflect downstream consequences, limiting their predictive value. Thus, these findings establish antigen-independent tonic signaling as a mechanistic driver of CAR-T efficacy and a clinically relevant marker to inform CAR design and improve therapeutic outcomes.

By defining a tonic signaling gene signature, we suggest a biomarker-guided framework for rational CAR optimization that expands beyond target selection to tuning basal signaling strength. Importantly, our results highlight that restraining, rather than enhancing ^69,70^, basal CAR activity preserves stemness, mitigates exhaustion, and enhances durability in DIPG/DMG. Thus, our studies reveal that tonic signaling may serve as a unifying axis that links CAR structure to T-cell function, persistence, and therapeutic efficacy. These insights shift the framework of CAR engineering and establish tonic signaling modulation as a potential strategy for achieving sustained efficacy against otherwise intractable solid tumors such as DIPG/DMG.

## Acknowledgements

We are grateful to Dr. Michelle Monje at the Stanford University for providing the DIPG cell lines. We thank Eva Nicholson for technical assistance and Dr. Ed Hurlock for his comments. This study was in part funded by grants from NIH (R01NS140460 and R01NS142389), the CancerFree KIDS, CureStartsNow Foundation, and ChadTough Defeat DIPG Foundation to Q.R.L.

## Author contributions

Conception and design: E.B.D., X.Z., D.X., and Q.R.L. Data acquisition, analysis, and interpretation: E.B.D., X.Z., D.X., U.K.S., W.M., X.H., M.C., P.-Y.L., J.B., Q.Q., S.M., M.H., A.E.B. Resources, writing and review: E.B.D., J.D., M.X., N.P.-S., H.R.T., C.B.S., J.B.F., P.D.B., S.R., C.K., J.A.C., Y.Z., and Q.R.L. Study supervision: Q.R.L.

## Competing interests

Authors declare that they have no competing interests.

## Data and materials availability

The high-throughput sequencing data that support the findings of this study have been deposited in the NCBI Gene Expression Omnibus (GEO) under accession codes GSE302513, GSE302514 and GSE305987.

## MATERIALS AND METHODS

### Cell lines and animals

The DIPG cell lines were cultured in tumor stem medium (TSM), prepared using a 1:1 mixture of Neurobasal medium and DMEM/F-12, supplemented with B-27 (without vitamin A), human EGF, human FGF, human PDGF-AA, human PDGF-BB, and heparin. All cell lines were maintained under standard neurosphere culture conditions. Immunodeficient NOD SCID gamma (NSG) mice were used for intracranial xenograft models. All animal use and study protocols were approved by the Institutional Animal Care and Use Committee at the Cincinnati Children′s Hospital Medical Center. Patient-derived DIPG cells were obtained and established with informed consent and in accordance with protocols approved by the Institutional Review Board (IRB) at Cincinnati Children’s Hospital Medical Center.

### Construction of chimeric antigen receptor (CAR) vectors

The single-chain variable fragment (scFv) sequences were derived from anti-B7-H3 monoclonal antibodies 376.96, MGA271, and Hu8H9 with human codon optimization. Each CAR construct consisted of a signal peptide (SP), scFv, hinge region, CD8 transmembrane domain (CD8TM), 4-1BB costimulatory domain, and CD3ζ signaling domain. The CAR sequence was linked to GFP via an internal ribosome entry site (IRES) and placed under the control of the elongation factor-1 alpha (EF1α) promoter. A mock control vector encoding only GFP under the EF1α promoter was used as a negative control. All constructs were synthesized and subcloned into a lentiviral backbone.

### T cell transduction and culture

Peripheral blood mononuclear cells (PBMCs) were isolated from healthy donors. CD4^+^ and CD8^+^ T cells were isolated from PBMC using REAlease® CD4/CD8 (TIL) MicroBead Kit (Miltenyi Biotec, #130-121-561) and activated using anti-CD3/CD28 Dynabeads (BioLegend, #422604). The isolated T cells were cultured in T cell medium consisting of RPMI 1640 supplemented with 10% fetal bovine serum (FBS), 5 ng/mL recombinant human IL-2 (MCE, Cat# HY-P7037), and 5 ng/mL recombinant human IL-15 (MCE, #HY-P7034). After 2 days of activation, T cells were infected with CAR-expressing lentivirus at a multiplicity of infection (MOI) of 5 in the presence of lentiboost (SIRION Biotech, #SB-P-LV-101-11) via spinoculation at 1500 × g for 90 minutes at 32°C. Transduced T cells were maintained and expanded in T cell medium under standard culture conditions.

### Flow cytometry

Cells were collected and analyzed using either the BD LSRFortessa™ or FACSymphony™ A5 cell analyzer (BD Biosciences) with the FACSDiVa software. For surface marker staining, T cells were incubated with antibodies for 30 min at 4°C in FACS buffer (PBS + 3% FBS). The fluorophore-conjugated anti-human antibodies used for surface staining were: CD3 APC (BioLegend, #300312), CD45RO APC (BioLegend, #304210), CD62L PE-Cy7 (BioLegend, #304822), CD45RA BV421 (BioLegend, #304129), CCR7 PerCP/Cy5.5 (BioLegend, #353219), PD-1 PE-Cy7 (BioLegend, #329918), TIM-3 PerCP/Cy5.5 (BioLegend, #345016), LAG-3 APC (BioLegend, #369212), CD25 BV421 (BioLegend, #302630), CD69 APC-Cy7 (BioLegend, #310914). For apoptosis analysis, cells were stained with Annexin V (BioLegend, #640932) according to the manufacturer’s protocol. For CAR-T affinity assays, the CAR-T cells were stained with different concentrations of AF647-conjugated Recombinant Human B7-H3 Protein (SinoBiological, #11188-H86H-SG) for 25 min at 4°C. For detecting the B7H3 expression on the membrane of DIPG cells, the cells were stained with APC-conjugated B7H3 antibody (BioLegend, #351006) for 30 min at 4°C. For proliferation assays, the T cells were stained with 5 μM ViaFluor® 405 SE dye (Biotium, #30068-T) for 15 min at room temperature according to the manufacturer’s protocol. The data was analyzed using FlowJo software. Sorting assays were performed using BD FACSymphony™ S6 cell sorter (BD Bioscience) by CCHMC Research Flow Cytometry Core (RFCC).

### CAR T cell imaging

CAR-T cells (5 × 10^5^) were stained with AF647-conjugated G4S linker antibody (CST, #69782S) on ice for 1 h. Cells were washed and resuspended in 200 ul buffer (PBS + 0.5% BSA). The cell suspension was added to the 8-well glass chamber for imaging. Images were acquired with confocal microscopy (Nikon) using a 20 × objective. Number of CAR clusters on the T cell membrane was quantified using ImageJ. To induce crosslinking of B7H3.BC CAR on the T cell membrane, the AF647-conjugated G4S linker antibody (CST, #69782S; 0.2 μg/ml) and goat anti-rabbit F(ab′)_2_ Fragment (Jackson ImmunoResearch, #111-006-047; 2.5 μg/ml) were added to the culture medium of B7H3.BC CAR-T cells on day 2 post-infection. The CAR-T cells were treated for 7-9 days, with antibody replenishment every 2 days.

### Enzyme-linked immunosorbent assay

To assess cytokine production by CAR-T cells, 5 × 10^4^ CAR-T cells were incubated with 5 × 10^4^ DIPG cells in 1 ml of DIPG cell culture medium in 24-well plates. After 24 hours of co-culture, cell-free supernatants were collected and analyzed for IFN-γ and IL-2 levels using commercially available ELISA kits (BioLegend), according to the manufacturer’s instructions. Each condition was assayed in duplicate.

### Western blot

Different B7-H3 CAR-T cells and control T cells were sorted for GFP⁺ cells on day 10 of culture. Total proteins were extracted from sorted and untransduced T cells using RIPA buffer supplemented with protease and phosphatase inhibitors. Protein samples were mixed with loading buffer, denatured at 100 °C for 10 minutes, and separated by 10% SDS-PAGE. Proteins were then transferred to 0.2 μm PVDF membranes (Bio-Rad, #1620177). After blocking with 5% nonfat milk in TBST for 1 hour at room temperature, membranes were incubated overnight at 4 °C with primary antibodies against STAT3 (CST, #12640S), phospho-STAT3 (CST, #9134P), AKT (CST, #9272S), phospho-AKT (CST, #9275S), S6 (CST, #2217S), phospho-S6 (CST, #2211S), and GAPDH (Thermo Fisher Scientific, #AM4300). The membranes were then incubated with HRP-conjugated secondary antibodies for 1 hour at room temperature. Protein bands were visualized using SuperSignalTM West Pico PLUS Chemiluminescent Substrate (Thermo Fisher Scientific, #34580).

### Luciferase assays

For luciferase assays, DIPG36 cells expressing luciferase were co-cultured with CAR-T cells or control T cells at effector (GFP+ T cells) to target (tumor cells) (E: T) ratio of 1:2 or 1:4 for two days. Then, the cells were lysed and the luciferase were detected using the Luciferase Assay System (Promega, #E4030) according to the manufacturer’s instructions.

### Co-culture imaging

In co-culture experiments using lentivirus-transduced CAR-T cells and DIPG cells, 5 × 10^4^ DIPG-C1, DIPG13, DIPG21 or DIPG36 cells expressing mcherry were co-cultured with lentivirus-infected CAR-T cells or control T cells (GFP+) at E:T ratio 1:2 or 1:4 in 8-well glass chambers for five days. The confocal images were taken using confocal microscopy (Nikon) at day 0, 1, 2, 3 and 5. Number of residual tumor cells was quantified using ImageJ. In co-culture experiments using lentivirus-transduced CAR-T cells and patient-derived DIPG spheroids, 5 × 10^4^ tumor cells were seeded in 8-well glass chambers 3 days to form the spheroids and then 2.5 × 10^4^ GFP+ CAR-T cells were added to the chamber. Confocal images were captured at indicated days post-CAR-T cell addition. In co-culture experiments using mRNA-based CAR-T cells and DIPG cells, 5 × 10^4^ DIPG13, DIPG21 or DIPG36 cells expressing mCherry were co-cultured with mRNA-based CAR-T cells or control T cells at E: T ratio of 1:1 in 8-well glass chambers for five days. The confocal images were taken at day 0, 1, 2, 3 and 5.

### 3D-tumor spheroid killing assay

To evaluate CAR-T cell-mediated cytotoxicity in a three-dimensional (3D) tumor model, DIPG36 spheroids were generated by seeding tumor cells into 96-well plates and incubating for 4-6 days to allow spheroid formation ^71^. Immediately after co-culture (E:T 1:10) with different CAR-T cells imaging at 0 hour was performed (as baseline) in optically compatible 8-well glass chamber. In 96-wells, the co-culture was then incubated for next 6 or 18 hours to enable effector-target interaction and killing. Before transferring for imaging, the egressed cells from 3D-tumor spheroids (due to CAR-T cell mediated killing) and unbound cellular debris were removed via gentle washing. The remaining intact or fragmented spheroids (engaged with CAR-T cells) were imaged by confocal microscopy. The tumor cell killing efficiency of both CAR-T cells were calculated by quantifying the mCherry fluorescence intensity of remaining tumor cells/fragments by ImageJ and after normalization with area average data was presented as bar graph.

### Rechallenge assays

To evaluate the long-term cytotoxic efficacy of CAR-T cells, a three-round sequential co-culture assay was performed. In the first round, mCherry-expressing DIPG13 cells were co-cultured with CAR-T cells or control T cells at an effector (GFP+ T cells) to target (DIPG13 cells) ratio of 1:4 for 3 days. Then, all the cells were collected and re-challenged with fresh DIPG13 cells for two additional rounds (Rounds 2 and 3). At the end of each round of co-culture, cells from duplicate wells were collected and analyzed by flow cytometry to determine the percentage of tumor cells (mCherry⁺) and T cells (CD3⁺). Absolute numbers of residual DIPG13 cells (mCherry⁺) and CAR-T cells (GFP⁺) were quantified using CountBright™ Absolute Counting Beads (Thermo Fisher Scientific, #C36950). Following the second round, the expression of exhaustion markers PD-1, LAG-3, and TIM-3 on CAR-T cells was evaluated by flow cytometry.

### Bulk RNA sequencing, CUT&Tag, and data analysis

Total RNA was extracted from sorted CAR-T cells and control T cells (GFP⁺), as well as untransduced T cells, on day 12 of culture using the RNeasy Mini Kit (Qiagen), followed by RNA sequencing. Differentially expressed genes were identified using the *limma* package in R (version 4.3.1), with thresholds set at |log_2_ fold change| > 2 and false discovery rate (FDR) < 0.05. Gene ontology (GO) enrichment analysis, principal component analysis (PCA), clustering analysis and gene expression heatmaps were generated using various R packages. Gene Set Enrichment Analysis (GSEA) was performed using gene sets obtained from the Molecular Signatures Database (MSigDB) and published literatures. Gene networks using selected pathways were identified using ToppCluster and visualized with Cytoscape (version 3.10.3). CUT & Tag assays were performed according to the manufacturer’s instructions using the sorted cells on day 12 of culture. The data were normalized across all samples using peak values near the transcription start sites (TSS) of several housekeeping genes as internal references. Resulting heatmaps were generated using deepTools, and genomic tracks were visualized with Integrative Genomics Viewer (IGV).

### Single-cell RNA sequencing (scRNA-seq)

Single-cell RNA sequencing was performed using the sorted CAR-T cells and control T cells (GFP⁺) cells on day 12 of culture. After sorting, dead cells were removed using a Dead Cell Removal kit (Miltenyi Biotec, #130-090-101). The cells were then resuspended using PBS with 0.04% BSA. Nearly 20,000–30,000 cells per sample were processed at the Cincinnati Children’s Hospital Medical Center (CCHMC) Single Cell Genomics Facility (RRID:SCR_022653) using the Chromium GEM-X Single Cell 3′ Kit v4 (Dual Index) on the Chromium X instrument (10x Genomics), according to the manufacturer’s protocol. The quality and concentration of the libraries were evaluated using the Agilent 2100 Bioanalyzer system, and sequencing was performed on an Illumina NovaSeq X Plus platform.

### scRNA-seq data analysis

Single-cell RNA sequencing (scRNA-seq) raw data were aligned to the human reference genome (GRCh38). The raw count matrix and BAM files were generated using Cell Ranger v9.0. Nuclear fraction scores were computed using the nuclear_fraction_annotation function in the DropletQC package, and cells with low nuclear fraction values were excluded. The raw count matrix was subsequently converted into a Seurat object using the Seurat v5.0 package. High-quality cells were defined as those expressing more than 500 and fewer than 7,500 genes, containing more than 3,000 unique molecular identifiers (UMIs), and exhibiting a mitochondrial transcript ratio below 15%. Data normalization was performed using the NormalizeData function, followed by principal component analysis (PCA) on the scaled expression matrix of the top 2,000 variable features identified by the FindVariableFeatures function. Batch effects were corrected using the Harmony package. Cell clustering and Uniform Manifold Approximation and Projection (UMAP) were performed using the top 25 Harmony components.

The cell clusters were annotated into nine T cell types: CD8 TSCM (CD8 stem cell memory T cells, *CD8A*^+^*KLF2*^high^*SELL*^high^), CD8 TCM (CD8 central memory T cells, *CD8A*^+^*KLF2*^medium^*SELL*^medium^*MKI67*^high^), CD8 TEM/TEFF (CD8 effector memory /effector T cells, *CD8A*^+^*KLF2*^low^*SELL*^low^*MKI67*^high^*GZMB*^high^), CD8 Tpex-like (CD8 progenitor exhausted T (Tpex)-like cells, *CD8A*^+^*KLF2*^low^*SELL*^low^*TCF7*^high^*CCR7*^high^*TOX2*^high^), CD8 TEX (CD8 exhausted T cells, *CD8A*^+^*KLF2*^low^*SELL*^low^*LAG3*^high^), CD4 TSCM/CM (CD4 stem cell memory / central memory T cells, *CD4*^+^*KLF2*^medium^*SELL*^medium^*IL7R*^high^), CD4 TEM (CD4 effector memory T cells, *CD4*^+^*KLF2*^low^*SELL*^low^*MKI67*^high^), CD4 Tpex-like (CD4 progenitor exhausted T (Tpex) -like cells, *CD4*^+^*KLF2*^low^*SELL*^low^*TCF7*^high^*CCR7*^high^*TOX2*^high^), CD4 TEFF/ TEX (CD4 effector / exhausted T cells, *CD4*^+^*KLF2*^low^*SELL*^low^*GZMA*^high^*HAVCR2*^high^). The AddModuleScore function was used to calculate the scores for gene signatures (eg. CAR-Ton, CAR-Tem, T cell exhaustion). scRNA-seq data were visualized using Seurat tools and R packages. Comparisons of gene set scores for multiple groups were performed using Kruskal-Wallis test followed by pairwise Wilcoxon rank-sum tests (BH-adjusted).

### CAR-T score analysis

The scRNA-seq dataset of GD2 CAR-T cells in DIPG was processed using the above pipeline, and the CAR-Ton score and T cell exhaustion score were calculated using the AddModuleScore function of Seurat. Comparisons of gene set scores between the two groups were performed using Wilcoxon rank-sum test. Three other published scRNA-seq datasets of CAR-T cells from clinical studies were also used for validation: Dataset 1 (*Deng et al., 2020*), Dataset 2 (*Haradhvala et al., 2022*), and Dataset 3 (*Steffin et al., 2024*). The scRNA-seq data were converted to pseudo-bulk RNA-seq data by calculating the average gene expression in CAR+ cells for each patient. Single-sample gene set enrichment analysis (ssGSEA; https://github.com/broadinstitute/ssGSEA2.0) were performed to get the scores using the GSVA package in R. Gene expression values were normalized to z-scores. Samples were stratified by clinical response into responder and non-responder groups based on the original study annotations. Comparisons of gene set scores between groups were performed using unpaired two-tailed Mann–Whitney test. Correlation heatmap and ROC curve analysis were conducted using the corresponding packages in R.

### Structure prediction for scFv dimers

The dimer structures of scFv were predicted using AlphaFold2-Multimer ^32,33^, with structural templates retrieved from the PDB100 database (version 20230517) included as part of the model input. After the initial predictions, a fast relaxation was performed using the AMBER force field ^72^. AlphaFold2-Multimer generates five independent dimer models, each accompanied by a confidence score. For each scFv sequence, the confidence scores across the five models were similar, indicating consistent structural predictions. Therefore, all five predicted structures were included in the subsequent binding affinity analysis.

### Binding affinity analysis using Rosetta scoring function

To ensure compatibility of the AlphaFold2-Multimer predicted structure with the Rosetta scoring function ^73^, we performed a fast relaxation using the same scoring method. Following relaxation, binding affinity was calculated as the difference between the score of the entire dimer complex and the sum of the scores of the individual protein structures.

### Steered molecular dynamics (SMD) simulation and free energy calculation

Molecular dynamics simulations were performed using GROMACS ^74^. The systems were first subjected to energy minimization using the steepest descent algorithm. Each system was built in a 7 × 10 × 7 nm water box containing 0.1 M NaCl. After minimization, the systems were equilibrated for 1 ns under NVT ensemble (constant number of particles, volume, and temperature) to stabilize the system at the target temperature and 1 ns under NPT ensemble (constant number of particles, pressure, and temperature) to adjust and reach equilibrium at the desired pressure conditions at 295 K and 1 bar. A 100 ns conventional MD simulation was then performed to further relax the complexes. Temperature and pressure were controlled using the Nosé–Hoover thermostat ^75^ and Parrinello–Rahman barostat ^76^, with time constants of 1 ps and 2 ps, respectively. All hydrogen-containing bonds were constrained using the LINCS algorithm ^77^. Following equilibration, SMD simulations were performed to pull one protein away from its interacting subunit. The last frame from the MD simulation was used as the starting structure. Due to differences in predicted binding poses, system dimensions were adjusted for each model. After adjustment, energy minimization and equilibration were repeated. Pulling was performed at a constant velocity of 9 nm/ns using a spring constant of 650 kJ/mol/nm^2^ ^74^. For each complex, 20 independent pulling simulations were conducted. Binding free energy was estimated using Jarzynski’s Equality based on the work profiles from these simulations ^35,36^.

### Xenograft mouse models

Immunodeficient NOD SCID gamma (NSG) mice were used for DIPG36 intracranial xenograft models. 8-week-old NSG mice were anesthetized and injected with 1 × 10^5^ luciferase-expressing DIPG36 cells into the pons regions. Three days post-tumor implantation, tumor burden was monitored by bioluminescence imaging using an IVIS Spectrum system. Then, the mice were randomized into individual groups and 1 × 10⁷ mRNA-electroporated CAR-T cells or control T cells were administered via intravenous (i.v.) injection. After CAR-T infusion, tumor burden was monitored weekly using IVIS. At experimental endpoints, mice were euthanized, and brains were harvested for histological analysis. Hematoxylin and eosin (H&E) staining was performed on paraffin-embedded brain sections to assess tumor infiltration and tissue morphology. For the DIPG36 orthotopic xenograft model, NSG mice were intracranially injected into the pons on day 0 with 1 × 10^5^ luciferase-expressing DIPG36 tumor cells along with 2 × 10^5^ lentivirus-transduced B7H3.BC CAR-T cells or control T cells. Tumor burden was monitored weekly using IVIS system. For the in vivo efficacy comparison of B7H3.BC CAR-T and MGA271 CAR-T, NSG mice were intracranially injected with 1 × 10^5^ luciferase-expressing DIPG36 tumor cells on day 0. After three days, 2.5 × 10^6^ B7H3.BC CAR-T, MGA271 CAR-T, or control T cells were injected into the tumor site. Tumor burden was monitored weekly using IVIS system. The animal studies were approved by the Institutional Animal Care and Use Committee of the Cincinnati Children’s Hospital Medical Center.

### Statistical analyses

Statistical analyses were performed using GraphPad Prism 8. Data were presented as dot plots or bar graphs showing the mean ± standard error of the mean (SEM) unless otherwise noted. *P* < 0.05 was considered statistically significant. Comparisons between two groups were conducted using two-tailed unpaired Student’s *t*-tests. Survival curves were analyzed using the log-rank test. Additional statistical methods were applied as indicated in the figure legends.

## Supplementary Figure Legends

**Figure S1.**
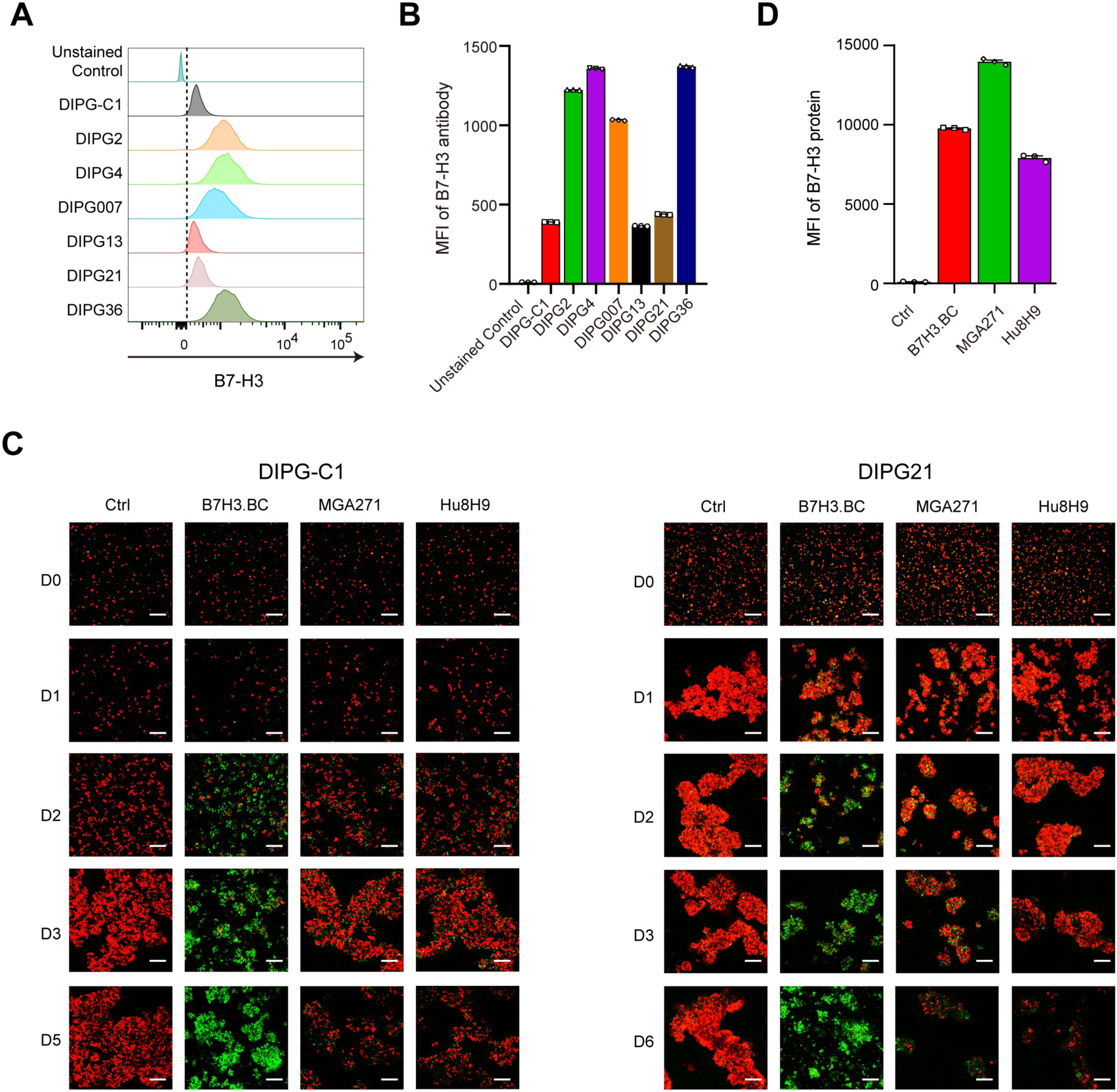
B7H3.BC CAR-T cells have better killing efficacy against DIPG cells in vitro than do MGA271 and Hu8H9 CAR-T cells. **(A)** Flow cytometric analysis of B7-H3 expression on DIPG cells. **(B)** Quantification of B7-H3 expression as measured by mean fluorescence intensity (MFI). Representative of three independent experiments. **(C)** Representative fluorescence images of B7-H3 or control CAR-T cells in co-culture experiments with DIPG-C1 (left) and DIPG21 (right) cells. The CAR-T cells (green) were co-cultured with DIPG cells (red) at a ratio of 1:4 for DIPG-C1 and 1:2 for DIPG21. Images were captured on days 0, 1, 2, 3, and 5 or 6. Representative of three donors. **(D)** MFIs of AF647-conjugated B7-H3 protein in cultures of indicated CAR-T cells. Representative of three independent experiments.

**Figure S2.**
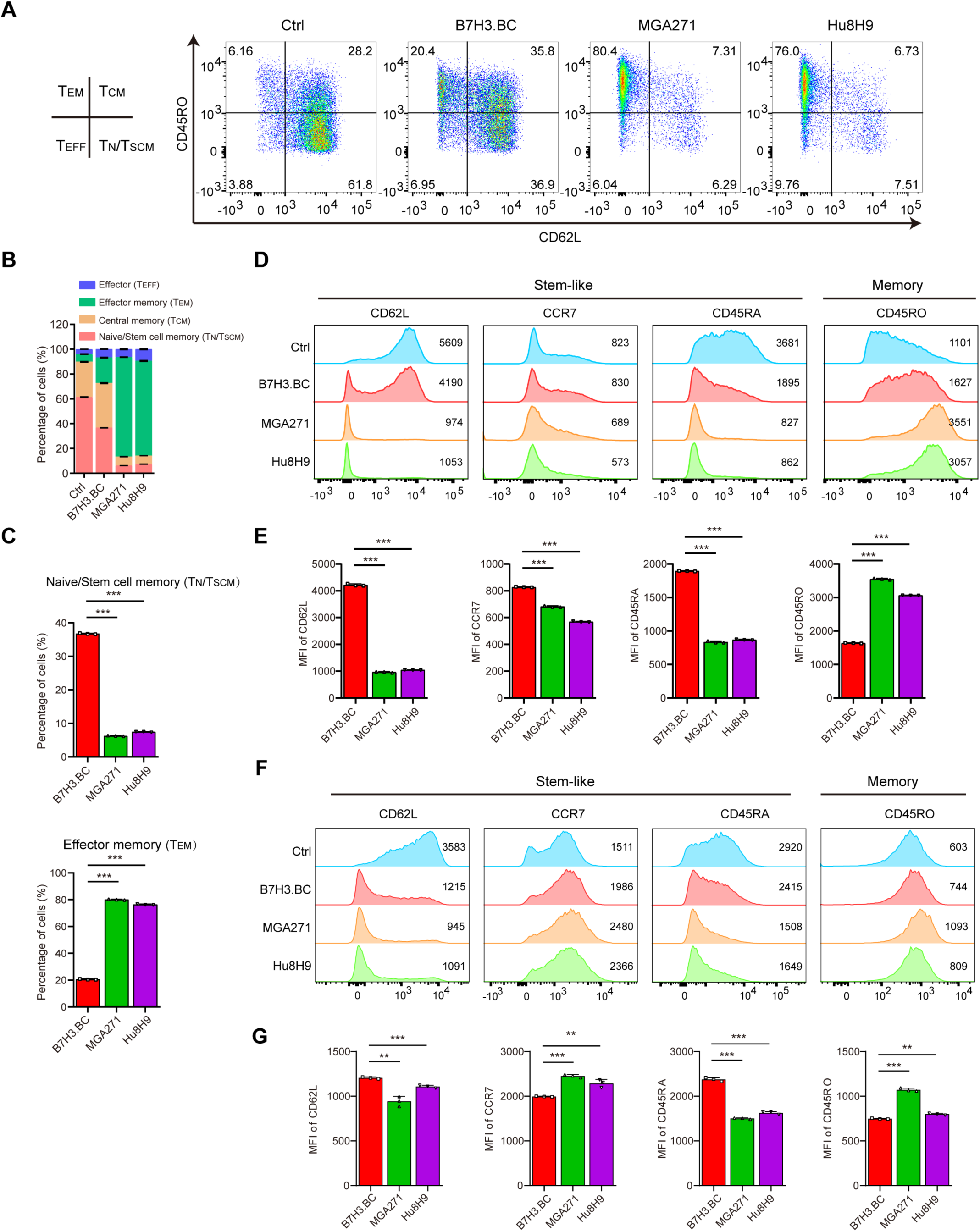
B7H3.BC CAR-T cells maintain a stem-like, less differentiated phenotype. **(A)** Representative flow cytometry plots showing the differentiation states of control and B7-H3 CAR-T cells on day 14 of culture. Surface expression of CD62L and CD45RO defines the following populations: naïve T cells/stem cell memory T cells (CD62L⁺CD45RO⁻), central memory T cells (CD62L⁺CD45RO⁺), effector memory T cells (CD62L⁻CD45RO⁺), and effector T cells (CD62L⁻CD45RO⁻). Representative of three donors. **(B)** Percentages of different populations in cultures of control and B7-H3 CAR-T cells on day 14 of culture. **(C)** Percentages of naïve T cells/stem cell memory T cells (left) and effector memory T cells (right) in cultures of control and B7-H3 CAR-T cells on day 14 of culture. **(D)** Representative flow cytometric analysis of stem-like markers (CD62L, CCR7, and CD45RA) and the memory marker (CD45RO) on CAR-T cells and control T cells after 14 days of culture. Representative of three donors. **(E)** Quantification of marker expression based on mean fluorescence intensity (MFI) CAR-T cells and control T cells after 14 days of culture. **(F)** Flow cytometric analysis of stemness markers CD62L, CCR7, and CD45RA and the memory marker CD45RO on CAR-T cells and control T cells after 7 days of culture. Representative of three donors. **(G)** Quantification of marker expression in panel based on MFI. For panels **C**, **E** and **G**, unpaired two-tailed Student’s *t*-test. **P < 0.01, ***P < 0.001.

**Figure S3.**
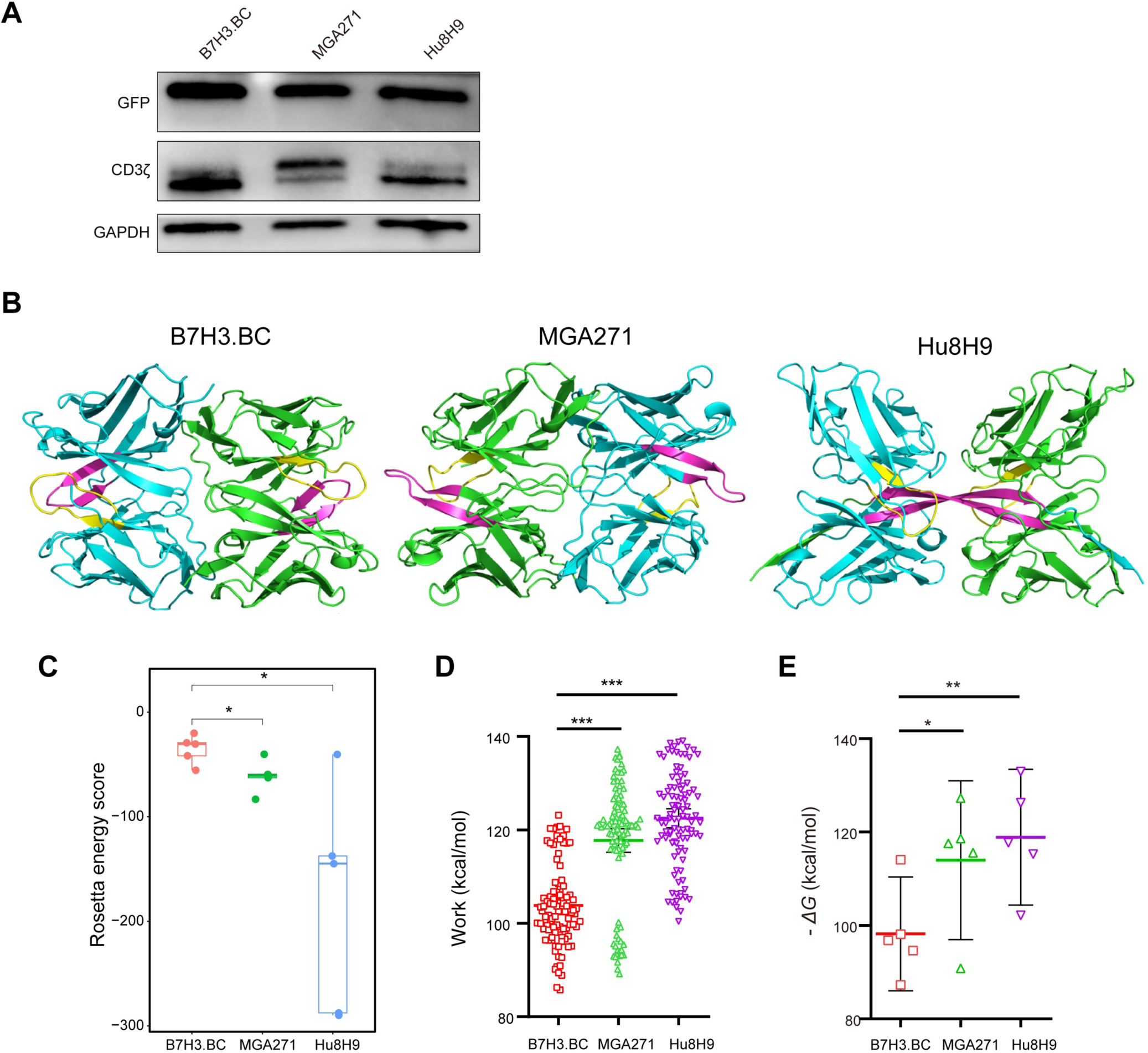
Detection of CAR expression and Structural modeling of scFv dimers and binding affinity analysis. **(A)** Western blot analysis for GFP and CAR CD3ζ in three B7-H3 CAR-T cells on day 10 of culture. **(B)** Representative dimer structures of each scFv predicted by AlphaFold2-Multimer, selected based on the highest confidence scores. Individual monomers are shown in blue and green. Blue and green coloring identifies different monomers. **(C)** The Rosetta energy score showing the binding affinity of three scFv dimers. **(D)** Pulling work measured by steered molecular dynamics (SMD) simulations for dissociating scFv monomers. A total of 100 replicas (20 replicas for each of 5 structural models) were shown. Error bars represent 95% CI. **(E)** Binding energy estimated using Jarzynski’s Equality based on 20 independent replicas of the pulling simulations for each dimer structure. Error bars represent 95% CI. **C-E**, unpaired one-tailed or two-tailed Student’s t-test. *P < 0.05, **P < 0.01, ***P < 0.001.

**Figure S4.**
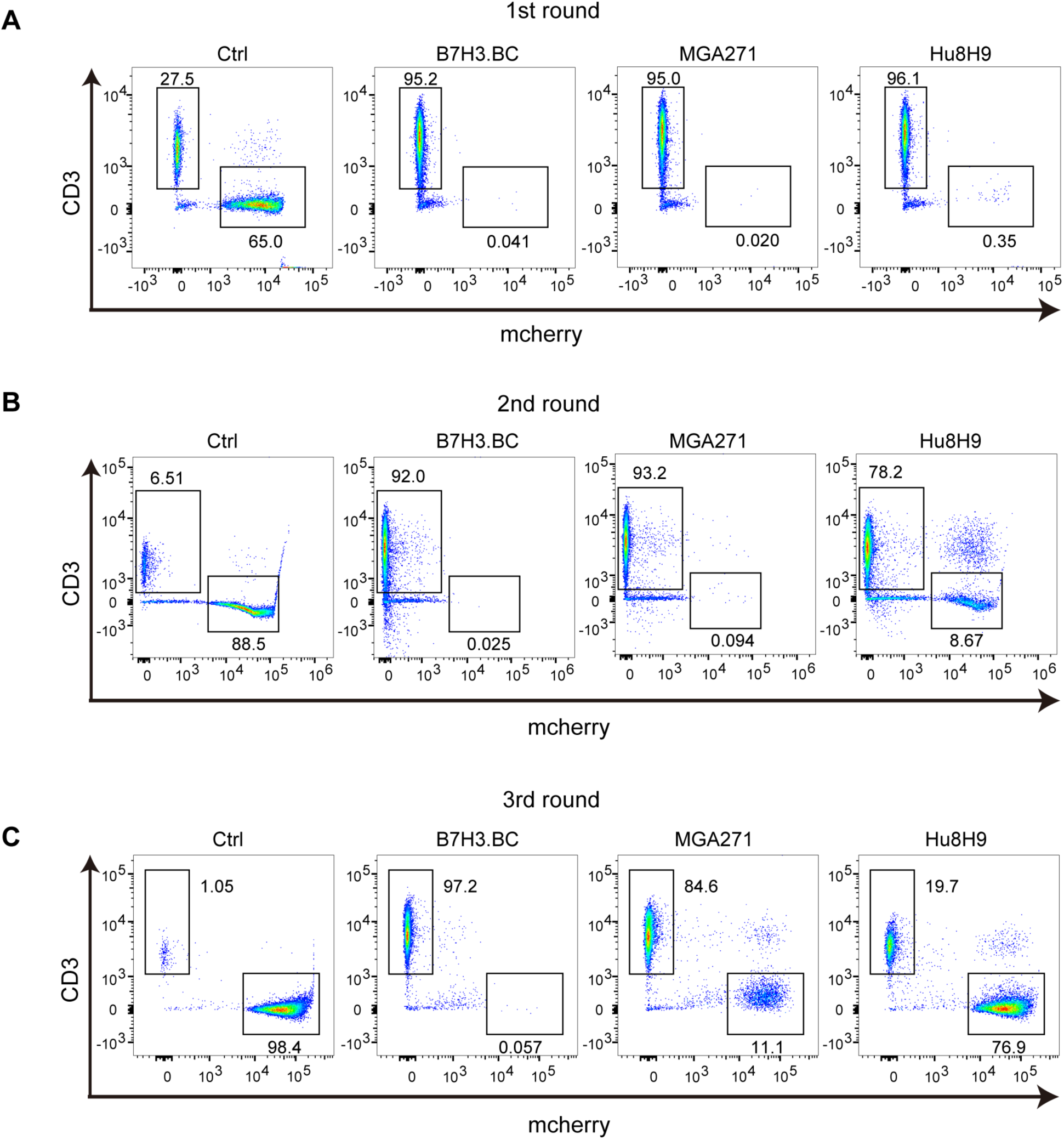
B7H3.BC CAR-T cells have long-term antitumor activity in rechallenge assays. **(A-C)** Representative flow cytometry plots of mCherry-labeled DIPG13 cells and GFP-labeled CAR-T cells after (**A**) one, (**B**) two, and (**C**) three rounds of co-culture. CAR-T cells and tumor cells were co-cultured at a ratio of 1:4. Representative of three donors.

**Figure S5.**
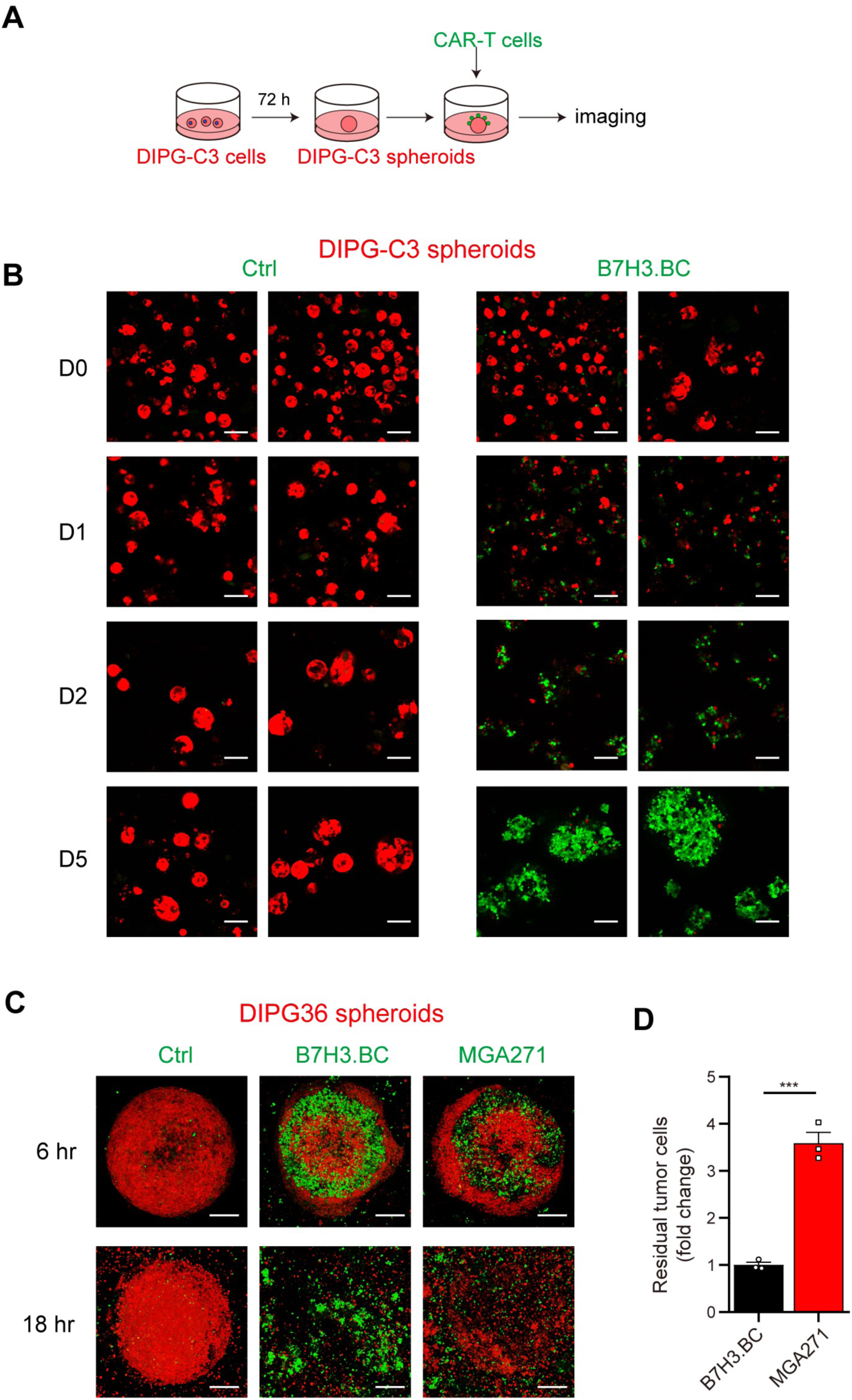
B7H3.BC CAR-T cells exhibit high efficacy in spheroid models. **(A)** Schematic of the co-culture experiment. Patient-derived DIPG cells were seeded in 8-well glass chambers to allow spheroid formation. After 3 days, CAR-T cells were added. **(B)** Representative fluorescence images of B7H3.BC CAR-T cells in co-culture with patient-derived DIPG spheroids. Images were captured at indicated days post-CAR-T cell addition. Representative of three donors. **(C)** Representative confocal images of B7H3.BC and MGA271 CAR-T cells in co-culture with DIPG36 3D-spheroids. Images were captured at indicated timepoints post-CAR-T cell addition. **(D)** mCherry intensities at 18 hours. Representative of three donors. **D**, unpaired two-tailed Student’s t-test. *P < 0.05.

**Figure S6.**
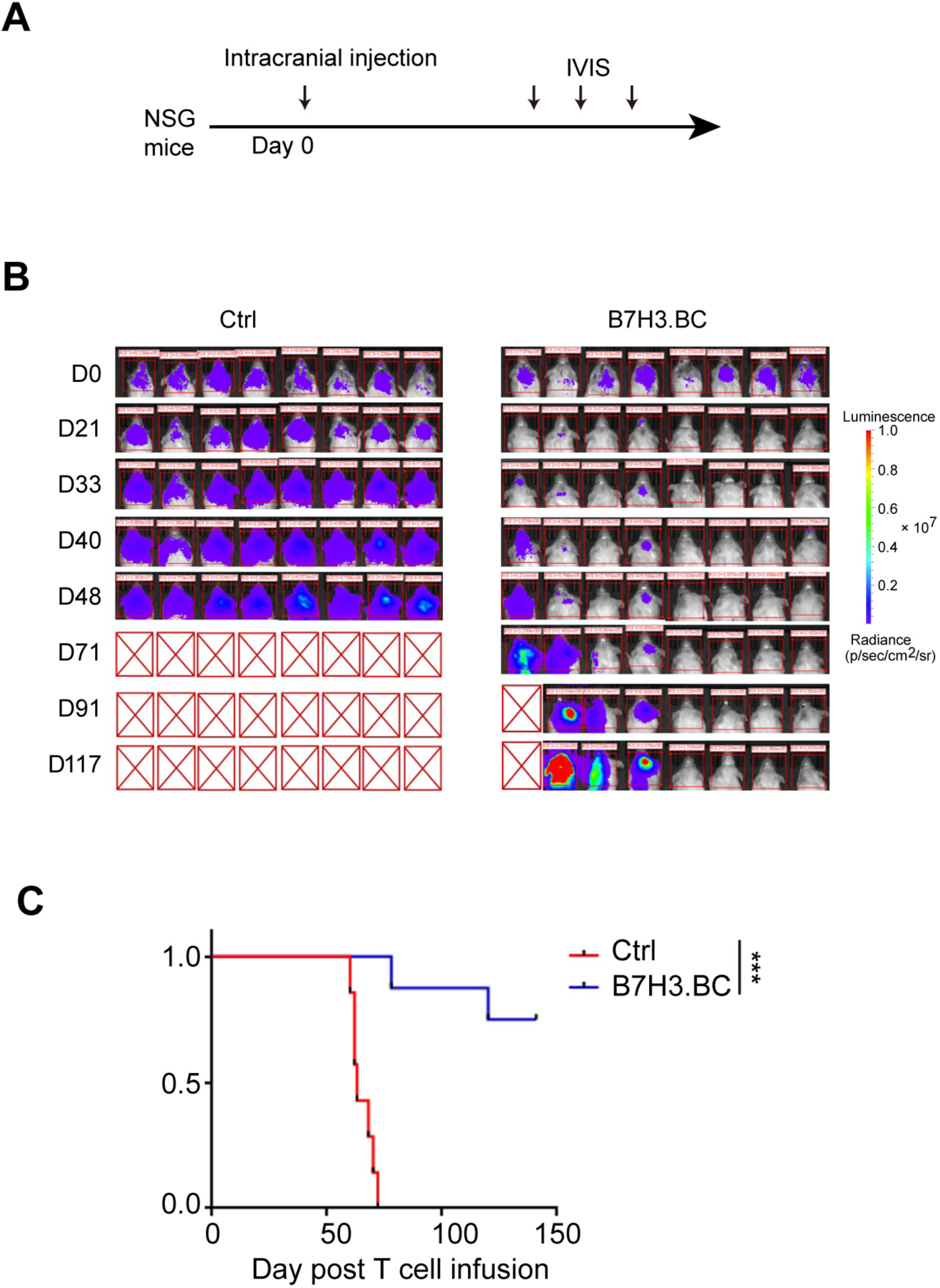
B7H3.BC CAR-T cells have high antitumor efficacy in vivo. **(A-C) A)** Schematic overview of the experiment. **B)** Bioluminescence images showing tumor progression (n = 8 per group). **C)** Kaplan-Meier survival curve of NSG mice (n = 8 per group).

**Figure S7.**
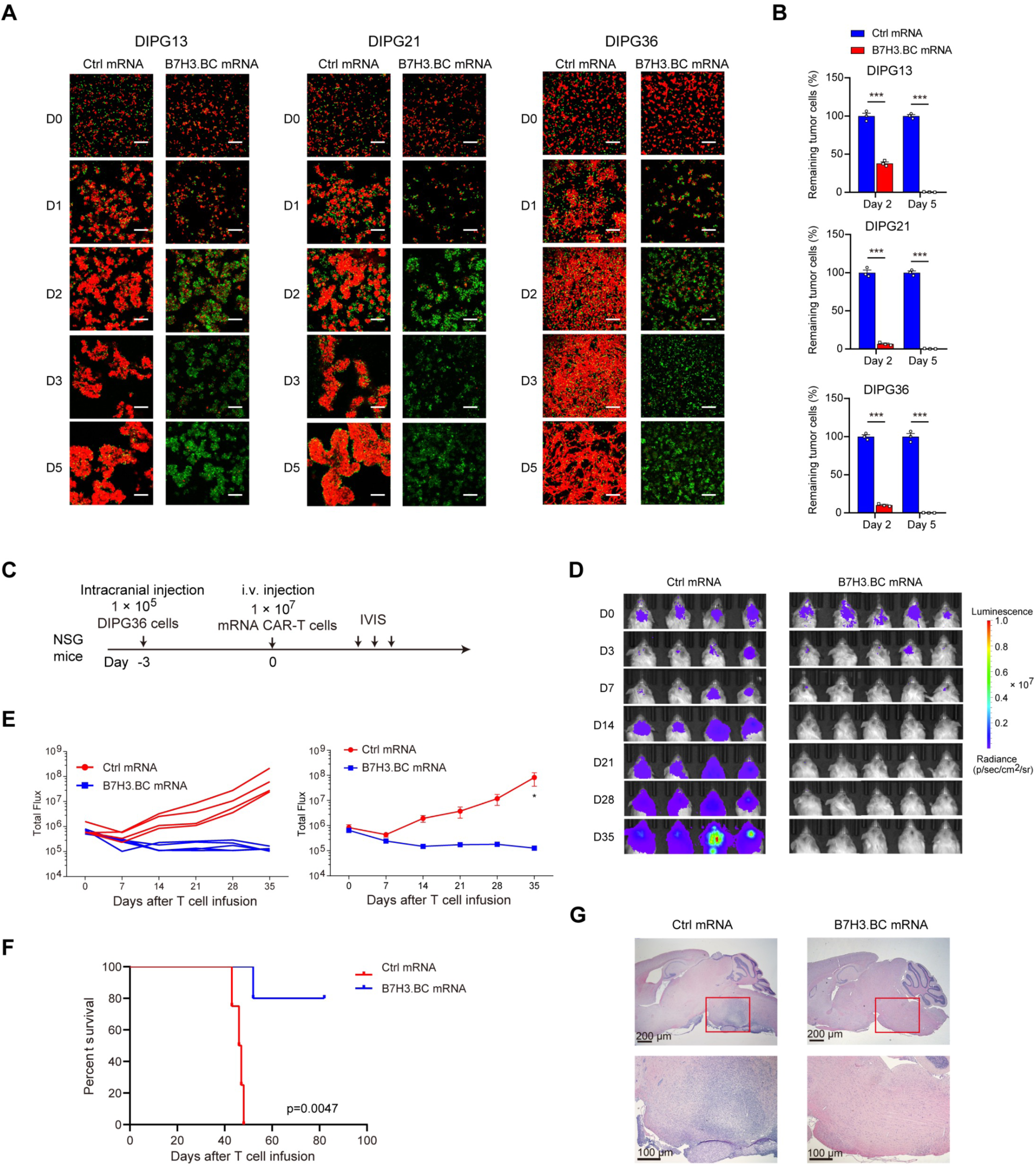
mRNA-based CAR-T cells demonstrate potent antitumor activity against DIPG. **(A)** Representative fluorescence images of B7H3.BC mRNA-based CAR-T cells in co-culture with indicated DIPG cells. The ratio of T cells (green) to DIPG cells (red) was 1:1. Scale bars = 200 µm. Representative of three donors. **(B)** Percentage survival of DIPG13 (upper), DIPG21 (middle) and DIPG36 (lower) at days 2 and 5 of incubation with indicated T cells. Data are means ± SEM. **C)** Schematic of experiment. **D)** Bioluminescence images of NSG mice. **E)** DIPG36 tumor growth over time. **F)** Kaplan-Meier survival curve. **G)** H&E staining of brain sections. The areas highlighted in red boxes are shown at higher magnification below. The dotted outlines indicate the boundaries of visible tumor regions. For panel **B**, *t*-test. ***P < 0.001. For panel **E**, unpaired Student’s *t*-test. For panel **F**, the log-rank test was used. *P < 0.05.

**Figure S8.**
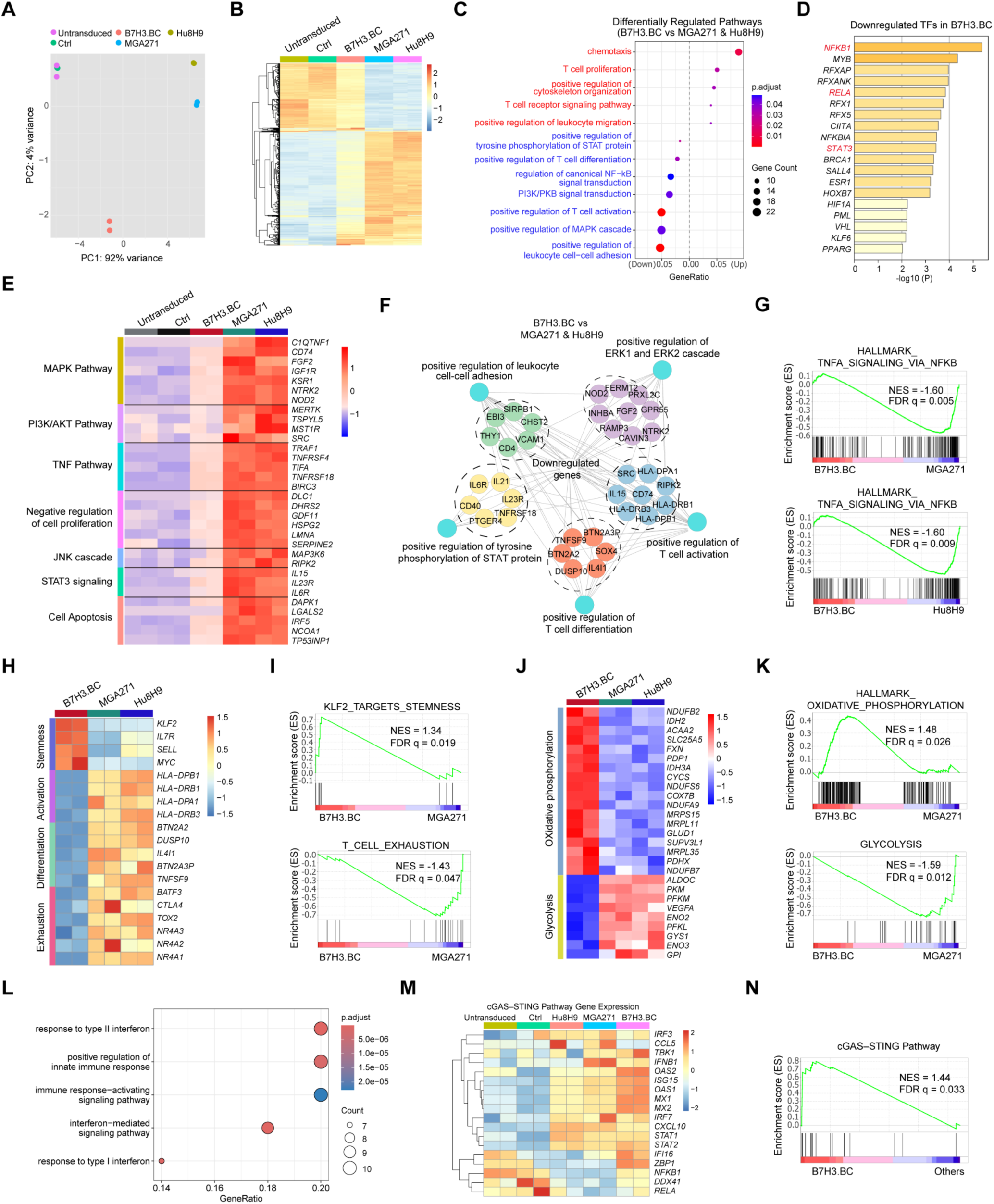
Transcriptomic analyses reveal a stem-like, less exhausted phenotype in B7H3.BC CAR-T. **(A)** Principal component analysis of RNA-seq data from untransduced T cells, control T cells (Ctrl), and B7H3.BC, MGA271 and Hu8H9 CAR-T cells on day 12 of culture. **(B)** Heatmap of differentially expressed genes across untransduced T cells, control T cells, and B7H3.BC, MGA271 and Hu8H9 CAR-T cells on day 12 of culture. **(C)** Gene Ontology analysis of upregulated pathways (red) and downregulated pathways (blue) in B7H3.BC CAR-T cells compared to MGA271 and Hu8H9 CAR-T cells. **(D)** Metascape transcription factor analysis of genes downregulated in B7H3.BC CAR-T cells compared to MGA271 and Hu8H9 CAR-T cells. **(E)** Heatmap of expression of selected genes associated with tonic signaling-related pathways. **(F)** Pathway enrichment analysis of genes downregulated in B7H3.BC CAR-T cells compared to MGA271 and Hu8H9 CAR-T cells. **(G)** Representative GSEA plots highlighting representative significantly downregulated pathways in B7H3.BC CAR-T cells compared to MGA271 or Hu8H9 CAR-T cells. **(H)** Heatmap of expression of selected genes related to stemness, activation, differentiation, and exhaustion across groups. **(I)** GSEA analysis showing differential enrichment of KLF2 target genes and the T cell exhaustion signature in B7H3.BC and MGA271 CAR-T cells. **(J)** Heatmap of expression of genes involved in oxidative phosphorylation and glycolysis pathways in B7H3.BC, MGA271, and Hu8H9 CAR-T cells relative to control T cells. **(K)** GSEA analysis highlighting representative metabolic pathways enriched in B7H3.BC CAR-T cells or in MGA271 and Hu8H9 CAR-T cells. **(L)** GO network analyses of pathways significantly upregulated in B7H3.BC CAR-T cells relative to the MGA271 and Hu8H9 CAR-T cells, control T cells transduced with empty vector (Ctrl), and untransduced T cells. **(M)** Heatmap of expression of genes in the cGAS-STING pathway across different groups. **(N)** GSEA analysis of the cGAS-STING pathway between B7H3.BC CAR-T and the other groups.

**Figure S9.**
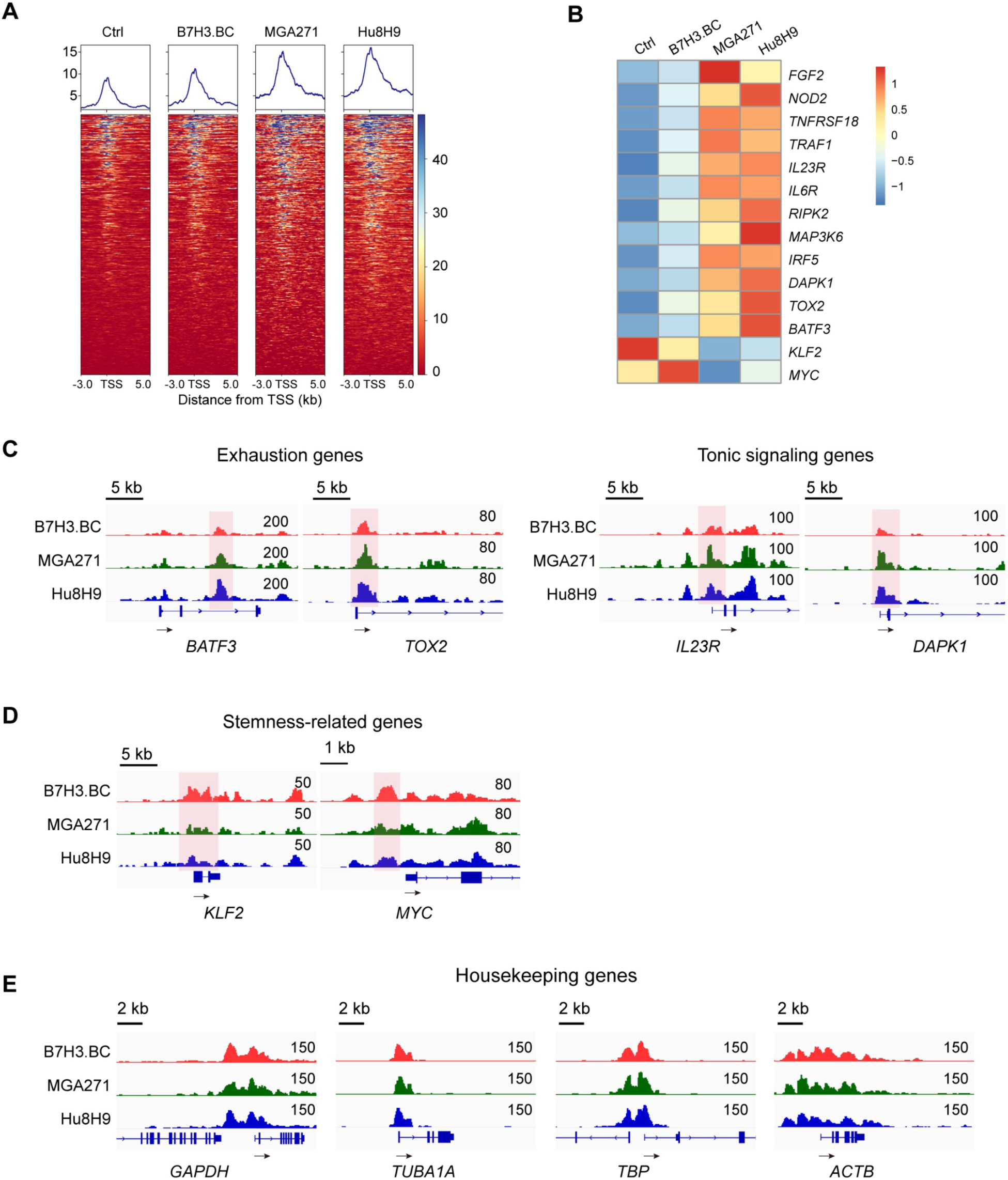
Epigenomic analyses reveal a stem-like, less exhausted phenotype. **(A)** Heatmap showing H3K27ac signal intensities near the TSS of genes downregulated in B7H3.BC CAR-T. The control T cells and CAR-T cells were cultured for 12 days after infection. **(B)** Heatmap of H3K27ac signal intensities at promoters or enhancers of associated with T cell exhaustion and tonic signaling in CAR-T cells relative to control T cells. **(C, D)** Representative tracks depicting H3K27ac enrichment at loci associated with **C)** T cell exhaustion, tonic signaling, and **D)** stemness-related genes in CAR-T cells. **(E)** Representative tracks depicting H3K27ac enrichment at the loci of housekeeping genes.

**Figure S10.**
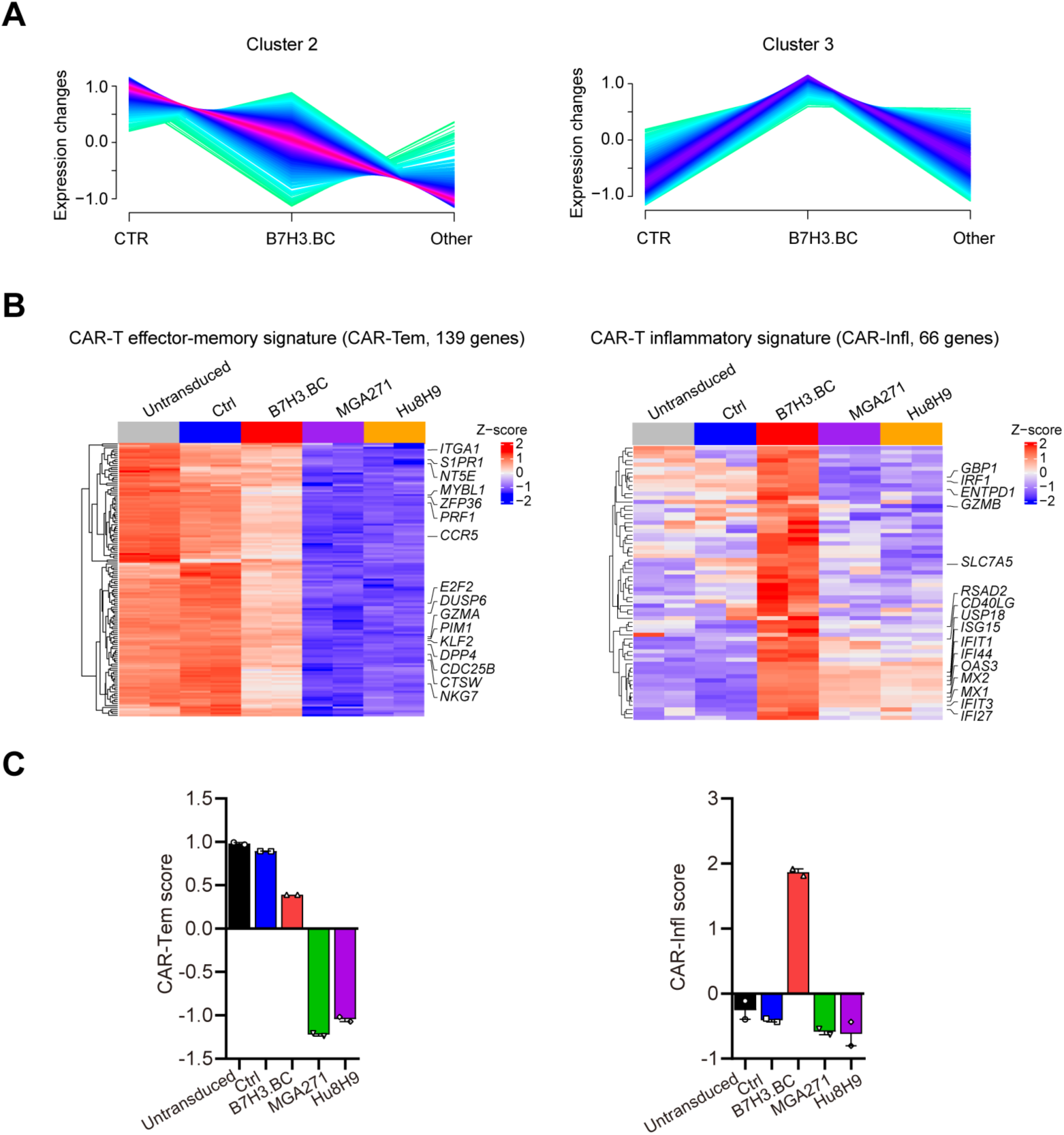
Clustering analysis of RNA-seq data to derive distinct CAR-T cell scores. **(A)** Clustering plots showing scaled expression changes of genes in CTR (untransduced and Ctrl), B7H3.BC CAR-T, and Other CAR-T (MGA271 and Hu8H9). Gene expression values were normalized to z-scores for visualization. **(B)** Heatmaps depicting differentially expressed genes associated with CAR-T effector-memory (CAR-Tem) signature, and CAR-T inflammatory (CAR-Infl) signature as indicated. **(C)** The CAR-Tem score and CAR-Infl score in untransduced T cells, control T cells, and B7H3.BC, MGA271 and Hu8H9 CAR-T cells. The scores were calculated by the ssGSEA method based on the corresponding gene lists.

**Figure S11.**
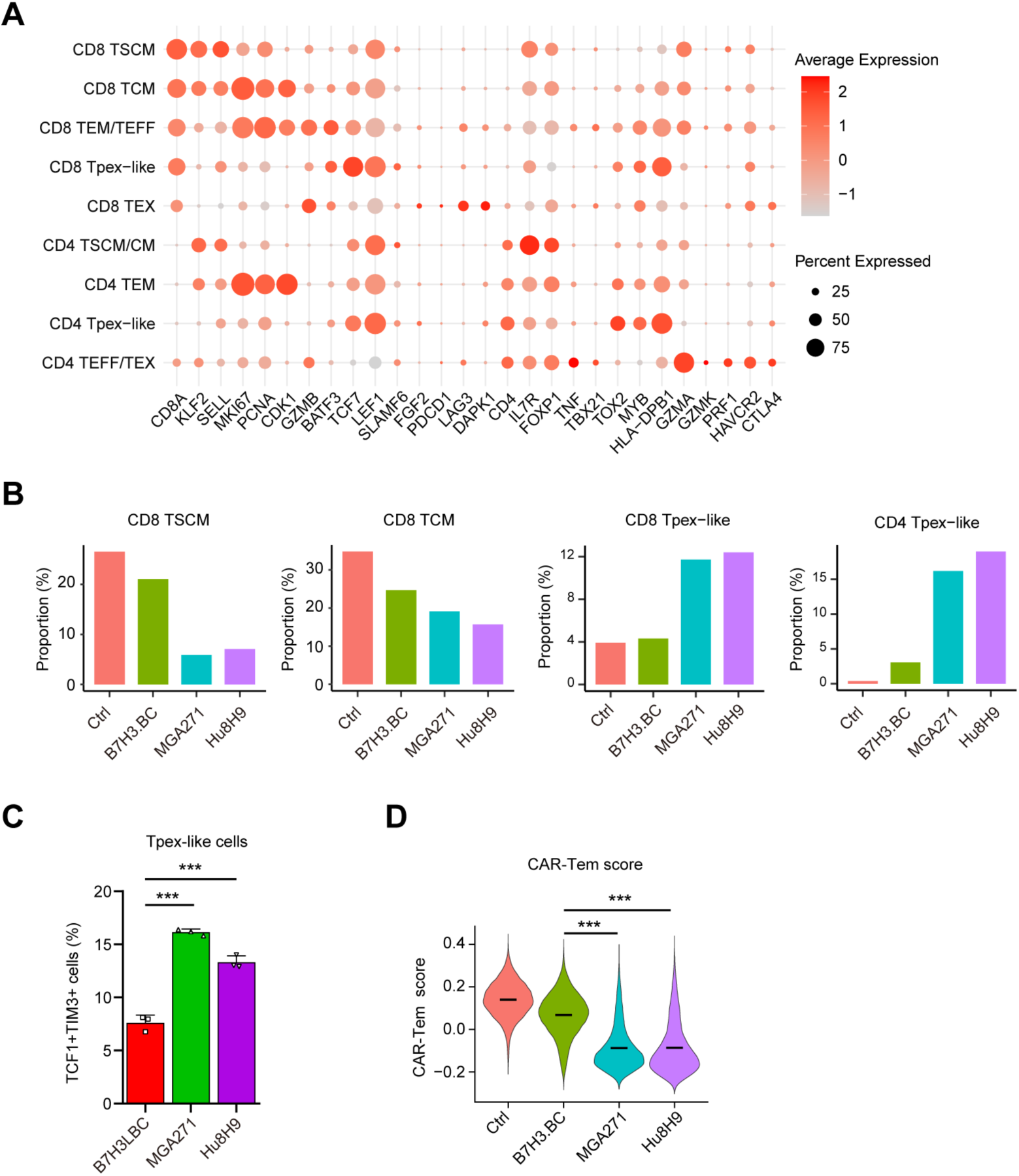
Single-cell RNA-seq analysis reveals changes across control and B7-H3 CAR-T cells. **(A)** Dot plot of marker gene expression in different T cell populations. Dot color represents the scaled average expression of marker genes in different cell populations. Dot size indicates the proportion of cells. **(B)** Proportions of CD8 TSCM, CD8 TCM and CD4/CD8 Tpex-like cells in control T and CAR-T cells. **(C)** Quantification of the proportion of TCF1^+^TIM3^+^ cells in B7-H3 CAR-T cells on day 13 of culture. Representative of three donors. **(D)** Violin plot of CAR-Tem scores in control and B7-H3 CAR-T cells. The crossbar represents mean value of the scores. **C**, unpaired two-tailed Student’s t-test. **D**, Kruskal-Wallis test followed by pairwise Wilcoxon rank-sum tests (BH-adjusted). ***P < 0.001.

**Figure S12.**
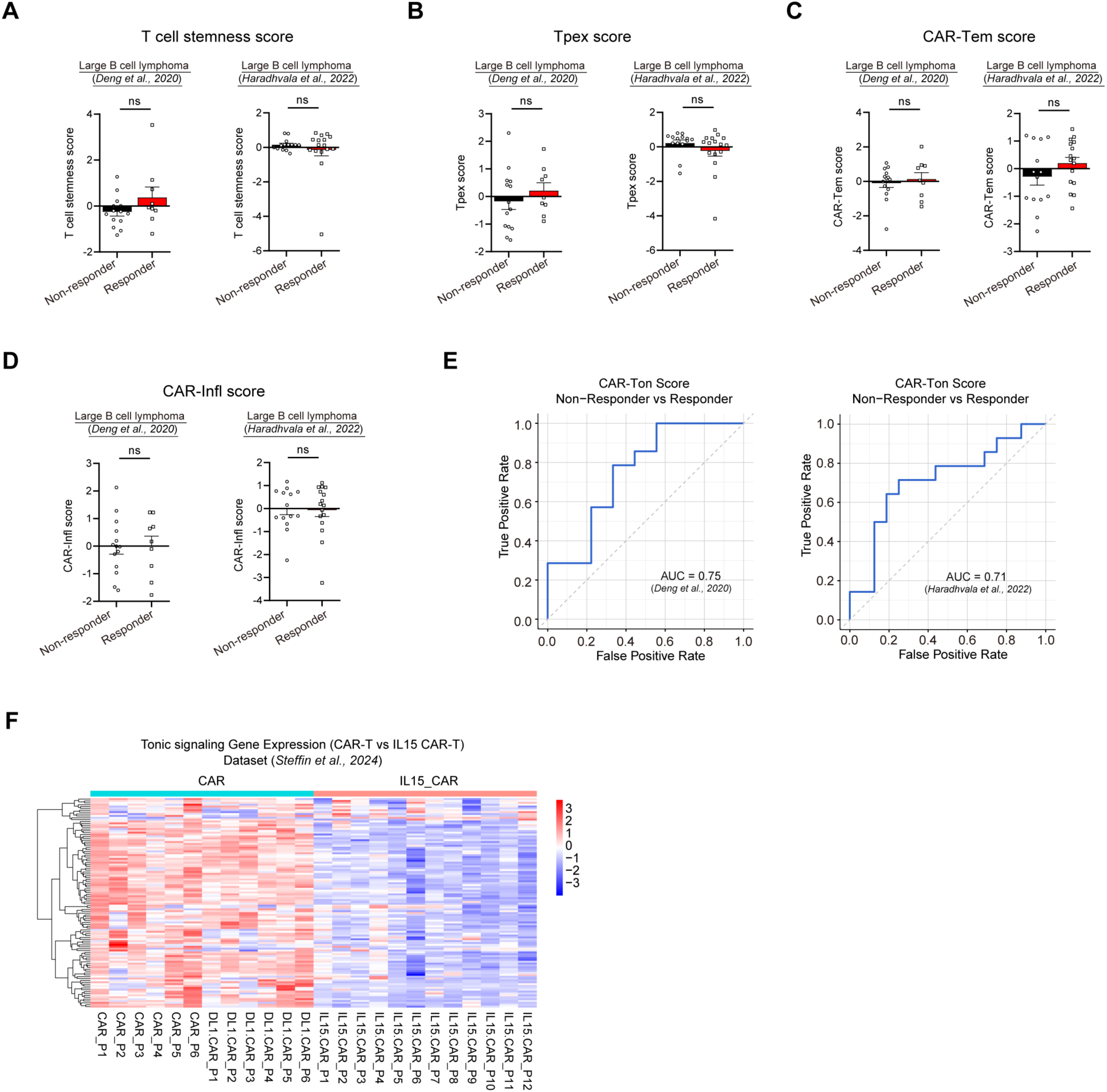
The correlation of CAR-T scores with clinical outcome. **(A-D)** Comparison of T cell stemness scores (**A**), Tpex scores (**B**), CAR-Tem scores (**C**) or CAR-Infl scores (**D**) in CAR-T non-responders and responders using the datasets from *Deng et al., 2020* (non-responder, n = 14; responder, n = 9) (left) or *Haradhvala et al., 2022* (non-responder, n = 14; responder, n = 16) (right). The scores were calculated using the ssGSEA method. **(E)** ROC curves with corresponding AUCs displaying the predictive performance of the CAR-Ton scores in distinguishing CAR-T non-responders from responders in the datasets from *Deng et al., 2020* (left) and *Haradhvala et al., 2022* (right), and calculated by the ssGSEA method. (**F**) Heatmap of tonic signaling gene expression in CAR-T and IL-15 CAR-T patients using the data from Dataset (*Steffin et al., 2024*). **A -D**, unpaired two-tailed Mann–Whitney test. ns, not significant.

## Notes

### Competing Interest Statement

The authors have declared no competing interest.

## References

1. Donaldson, S.S., Laningham, F., and Fisher, P.G. (2006). Advances toward an understanding of brainstem gliomas. J Clin Oncol 24, 1266–1272. 10.1200/JCO.2005.04.6599.

2. Hargrave, D., Bartels, U., and Bouffet, E. (2006). Diffuse brainstem glioma in children: critical review of clinical trials. Lancet Oncol 7, 241–248. 10.1016/S1470-2045(06)70615-5.

3. Johung, T.B., and Monje, M. (2017). Diffuse Intrinsic Pontine Glioma: New Pathophysiological Insights and Emerging Therapeutic Targets. Curr Neuropharmacol 15, 88–97. 10.2174/1570159x14666160509123229.

4. Marofi, F., Achmad, H., Bokov, D., Abdelbasset, W.K., Alsadoon, Z., Chupradit, S., Suksatan, W., Shariatzadeh, S., Hasanpoor, Z., Yazdanifar, M., et al. (2022). Hurdles to breakthrough in CAR T cell therapy of solid tumors. Stem Cell Res Ther 13, 140. 10.1186/s13287-022-02819-x.

5. Du, B., Qin, J., Lin, B., Zhang, J., Li, D., and Liu, M. (2025). CAR-T therapy in solid tumors. Cancer Cell 43, 665–679. 10.1016/j.ccell.2025.03.019.

6. Singh, N., and Maus, M.V. (2023). Synthetic manipulation of the cancer-immunity cycle: CAR-T cell therapy. Immunity 56, 2296–2310. 10.1016/j.immuni.2023.09.010.

7. Timpanaro, A., Song, E.Z., Amwas, N., Chiu, C.H., Ronsley, R., Taylor, M.R., Foster, J.B., Wang, L.D., and Vitanza, N.A. (2025). Evolving CAR T-Cell Therapy to Overcome the Barriers in Treating Pediatric Central Nervous System Tumors. Cancer Discov 15, 890–902. 10.1158/2159-8290.CD-24-1465.

8. Brown, C.E., Alizadeh, D., Starr, R., Weng, L., Wagner, J.R., Naranjo, A., Ostberg, J.R., Blanchard, M.S., Kilpatrick, J., Simpson, J., et al. (2016). Regression of Glioblastoma after Chimeric Antigen Receptor T-Cell Therapy. N Engl J Med 375, 2561–2569. 10.1056/NEJMoa1610497.

9. Majzner, R.G., Ramakrishna, S., Yeom, K.W., Patel, S., Chinnasamy, H., Schultz, L.M., Richards, R.M., Jiang, L., Barsan, V., Mancusi, R., et al. (2022). GD2-CAR T cell therapy for H3K27M-mutated diffuse midline gliomas. Nature 603, 934–941. 10.1038/s41586-022-04489-4.

10. Monje, M., Mahdi, J., Majzner, R., Yeom, K.W., Schultz, L.M., Richards, R.M., Barsan, V., Song, K.W., Kamens, J., Baggott, C., et al. (2025). Intravenous and intracranial GD2-CAR T cells for H3K27M(+) diffuse midline gliomas. Nature 637, 708–715. 10.1038/s41586-024-08171-9.

11. Thomas, B.C., Staudt, D.E., Douglas, A.M., Monje, M., Vitanza, N.A., and Dun, M.D. (2023). CAR T cell therapies for diffuse midline glioma. Trends Cancer 9, 791–804. 10.1016/j.trecan.2023.07.007.

12. Das, A.K., Sinha, M., Singh, S.K., Chaudhary, A., Boro, A.K., Agrawal, M., Bhardwaj, S., Kishore, S., and Kumari, K. (2024). CAR T-cell therapy: a potential treatment strategy for pediatric midline gliomas. Acta Neurol Belg 124, 1251–1261. 10.1007/s13760-024-02519-8.

13. Sterner, R.C., and Sterner, R.M. (2021). CAR-T cell therapy: current limitations and potential strategies. Blood Cancer J 11, 69. 10.1038/s41408-021-00459-7.

14. Levstek, L., Janzic, L., Ihan, A., and Kopitar, A.N. (2024). Biomarkers for prediction of CAR T therapy outcomes: current and future perspectives. Front Immunol 15, 1378944. 10.3389/fimmu.2024.1378944.

15. Xiong, D., Yu, H., and Sun, Z.J. (2024). Unlocking T cell exhaustion: Insights and implications for CAR-T cell therapy. Acta Pharm Sin B 14, 3416–3431. 10.1016/j.apsb.2024.04.022.

16. Hege, K. (2021). Context matters in CAR T cell tonic signaling. Nat Med 27, 763–764. 10.1038/s41591-021-01340-7.

17. Huang, Y., and Wang, H. (2025). Tonic signaling in CAR-T therapy: the lever long enough to move the planet. Front Med. 10.1007/s11684-025-1130-x.

18. Long, A.H., Haso, W.M., Shern, J.F., Wanhainen, K.M., Murgai, M., Ingaramo, M., Smith, J.P., Walker, A.J., Kohler, M.E., Venkateshwara, V.R., et al. (2015). 4-1BB costimulation ameliorates T cell exhaustion induced by tonic signaling of chimeric antigen receptors. Nat Med 21, 581–590. 10.1038/nm.3838.

19. Barden, M., Elsenbroich, P.R., Haas, V., Ertelt, M., Pervan, P., Velas, L., Gergely, B., Szoor, A., Harrer, D.C., Bezler, V., et al. (2024). Integrating binding affinity and tonic signaling enables a rational CAR design for augmented T cell function. J Immunother Cancer 12. 10.1136/jitc-2024-010208.

20. Ajina, A., and Maher, J. (2018). Strategies to Address Chimeric Antigen Receptor Tonic Signaling. Mol Cancer Ther 17, 1795–1815. 10.1158/1535-7163.MCT-17-1097.

21. Guedan, S., Madar, A., Casado-Medrano, V., Shaw, C., Wing, A., Liu, F., Young, R.M., June, C.H., and Posey, A.D., Jr. (2020). Single residue in CD28-costimulated CAR-T cells limits long-term persistence and antitumor durability. J Clin Invest 130, 3087–3097. 10.1172/JCI133215.

22. Zhou, Z., Luther, N., Ibrahim, G.M., Hawkins, C., Vibhakar, R., Handler, M.H., and Souweidane, M.M. (2013). B7-H3, a potential therapeutic target, is expressed in diffuse intrinsic pontine glioma. J Neurooncol 111, 257–264. 10.1007/s11060-012-1021-2.

23. Vitanza, N.A., Wilson, A.L., Huang, W., Seidel, K., Brown, C., Gustafson, J.A., Yokoyama, J.K., Johnson, A.J., Baxter, B.A., Koning, R.W., et al. (2023). Intraventricular B7-H3 CAR T Cells for Diffuse Intrinsic Pontine Glioma: Preliminary First-in-Human Bioactivity and Safety. Cancer Discov 13, 114–131. 10.1158/2159-8290.CD-22-0750.

24. Majzner, R.G., Theruvath, J.L., Nellan, A., Heitzeneder, S., Cui, Y., Mount, C.W., Rietberg, S.P., Linde, M.H., Xu, P., Rota, C., et al. (2019). CAR T Cells Targeting B7-H3, a Pan-Cancer Antigen, Demonstrate Potent Preclinical Activity Against Pediatric Solid Tumors and Brain Tumors. Clin Cancer Res 25, 2560–2574. 10.1158/1078-0432.CCR-18-0432.

25. Li, N., Zhang, C., Li, X., Liu, S., Xu, Y., and Yang, X. (2025). Targeting B7-H3 in solid tumors: Development and evaluation of novel CAR-T Cell therapy. Immunobiology 230, 152888. 10.1016/j.imbio.2025.152888.

26. Du, H., Hirabayashi, K., Ahn, S., Kren, N.P., Montgomery, S.A., Wang, X., Tiruthani, K., Mirlekar, B., Michaud, D., Greene, K., et al. (2019). Antitumor Responses in the Absence of Toxicity in Solid Tumors by Targeting B7-H3 via Chimeric Antigen Receptor T Cells. Cancer Cell 35, 221–237 e228. 10.1016/j.ccell.2019.01.002.

27. Vitanza, N.A., Ronsley, R., Choe, M., Seidel, K., Huang, W., Rawlings-Rhea, S.D., Beam, M., Steinmetzer, L., Wilson, A.L., Brown, C., et al. (2025). Intracerebroventricular B7-H3-targeting CAR T cells for diffuse intrinsic pontine glioma: a phase 1 trial. Nat Med 31, 861–868. 10.1038/s41591-024-03451-3.

28. Takeshita, M., Suzuki, K., Kassai, Y., Takiguchi, M., Nakayama, Y., Otomo, Y., Morita, R., Miyazaki, T., Yoshimura, A., and Takeuchi, T. (2015). Polarization diversity of human CD4+ stem cell memory T cells. Clin Immunol 159, 107–117. 10.1016/j.clim.2015.04.010.

29. Wang, X., Wong, C.W., Urak, R., Taus, E., Aguilar, B., Chang, W.C., Mardiros, A., Budde, L.E., Brown, C.E., Berger, C., et al. (2016). Comparison of naive and central memory derived CD8(+) effector cell engraftment fitness and function following adoptive transfer. Oncoimmunology 5, e1072671. 10.1080/2162402X.2015.1072671.

30. Guegan, J.P., Peyraud, F., Dadone-Montaudie, B., Teyssonneau, D., Palmieri, L.J., Clot, E., Cousin, S., Roubaud, G., Cabart, M., Leroy, L., et al. (2024). Analysis of PD1, LAG3, TIGIT, and TIM3 expression in human lung adenocarcinoma reveals a 25-gene signature predicting immunotherapy response. Cell Rep Med 5, 101831. 10.1016/j.xcrm.2024.101831.

31. Liu, Y., Liu, G., Wang, J., Zheng, Z.Y., Jia, L., Rui, W., Huang, D., Zhou, Z.X., Zhou, L., Wu, X., et al. (2021). Chimeric STAR receptors using TCR machinery mediate robust responses against solid tumors. Sci Transl Med 13. 10.1126/scitranslmed.abb5191.

32. Evans, R., O’Neill, M., Pritzel, A., Antropova, N., Senior, A., Green, T., Žídek, A., Bates, R., Blackwell, S., and Yim, J. (2021). Protein complex prediction with AlphaFold-Multimer. biorxiv, 2021.2010. 2004.463034.

33. Jumper, J., Evans, R., Pritzel, A., Green, T., Figurnov, M., Ronneberger, O., Tunyasuvunakool, K., Bates, R., Zidek, A., Potapenko, A., et al. (2021). Highly accurate protein structure prediction with AlphaFold. Nature 596, 583–589. 10.1038/s41586-021-03819-2.

34. Cheung, J., Wazir, S., Bell, D.R., Kochenderfer, J.N., Hendrickson, W.A., and Youkharibache, P. (2023). Crystal structure of a chimeric antigen receptor (CAR) scFv domain rearrangement forming a VL-VL dimer. Crystals 13, 710.

35. Jarzynski, C. (1997). Equilibrium free-energy differences from nonequilibrium measurements: A master-equation approach. Physical Review E 56, 5018.

36. Park, S., Khalili-Araghi, F., Tajkhorshid, E., and Schulten, K. (2003). Free energy calculation from steered molecular dynamics simulations using Jarzynski’s equality. The Journal of chemical physics 119, 3559–3566.

37. Wang, D., Starr, R., Alizadeh, D., Yang, X., Forman, S.J., and Brown, C.E. (2019). In Vitro Tumor Cell Rechallenge For Predictive Evaluation of Chimeric Antigen Receptor T Cell Antitumor Function. J Vis Exp. 10.3791/59275.

38. Prinzing, B., Zebley, C.C., Petersen, C.T., Fan, Y., Anido, A.A., Yi, Z., Nguyen, P., Houke, H., Bell, M., Haydar, D., et al. (2021). Deleting DNMT3A in CAR T cells prevents exhaustion and enhances antitumor activity. Sci Transl Med 13, eabh0272. 10.1126/scitranslmed.abh0272.

39. Dimitri, A.J., Baxter, A.E., Chen, G.M., Hopkins, C.R., Rouin, G.T., Huang, H., Kong, W., Holliday, C.H., Wiebking, V., Bartoszek, R., et al. (2024). TET2 regulates early and late transitions in exhausted CD8(+) T cell differentiation and limits CAR T cell function. Sci Adv 10, eadp9371. 10.1126/sciadv.adp9371.

40. Meister, H., Look, T., Roth, P., Pascolo, S., Sahin, U., Lee, S., Hale, B.D., Snijder, B., Regli, L., Ravi, V.M., et al. (2022). Multifunctional mRNA-Based CAR T Cells Display Promising Antitumor Activity Against Glioblastoma. Clin Cancer Res 28, 4747–4756. 10.1158/1078-0432.CCR-21-4384.

41. Qin, S., Tang, X., Chen, Y., Chen, K., Fan, N., Xiao, W., Zheng, Q., Li, G., Teng, Y., Wu, M., and Song, X. (2022). mRNA-based therapeutics: powerful and versatile tools to combat diseases. Signal Transduct Target Ther 7, 166. 10.1038/s41392-022-01007-w.

42. Foster, J.B., Madsen, P.J., Harvey, K., Griffin, C., Stern, A., Patterson, L., Joshi, N., Dickson, C., McManus, O., Beaubien, E., et al. (2025). Transient mRNA CAR T cells targeting GD2 provide dose-adjusted efficacy against diffuse midline glioma and high grade glioma models. Neuro Oncol. 10.1093/neuonc/noaf115.

43. Xia, A., Zhang, Y., Xu, J., Yin, T., and Lu, X.J. (2019). T Cell Dysfunction in Cancer Immunity and Immunotherapy. Front Immunol 10, 1719. 10.3389/fimmu.2019.01719.

44. Selli, M.E., Landmann, J.H., Terekhova, M., Lattin, J., Heard, A., Hsu, Y.S., Chang, T.C., Chang, J., Warrington, J., Ha, H., et al. (2023). Costimulatory domains direct distinct fates of CAR-driven T-cell dysfunction. Blood 141, 3153–3165. 10.1182/blood.2023020100.

45. Chan, J.D., Scheffler, C.M., Munoz, I., Sek, K., Lee, J.N., Huang, Y.K., Yap, K.M., Saw, N.Y.L., Li, J., Chen, A.X.Y., et al. (2024). FOXO1 enhances CAR T cell stemness, metabolic fitness and efficacy. Nature 629, 201–210. 10.1038/s41586-024-07242-1.

46. Patsoukis, N., Bardhan, K., Weaver, J., Herbel, C., Seth, P., Li, L., and Boussiotis, V.A. (2016). The role of metabolic reprogramming in T cell fate and function. Curr Trends Immunol 17, 1–12.

47. Peng, H.Y., Lucavs, J., Ballard, D., Das, J.K., Kumar, A., Wang, L., Ren, Y., Xiong, X., and Song, J. (2021). Metabolic Reprogramming and Reactive Oxygen Species in T Cell Immunity. Front Immunol 12, 652687. 10.3389/fimmu.2021.652687.

48. Creyghton, M.P., Cheng, A.W., Welstead, G.G., Kooistra, T., Carey, B.W., Steine, E.J., Hanna, J., Lodato, M.A., Frampton, G.M., Sharp, P.A., et al. (2010). Histone H3K27ac separates active from poised enhancers and predicts developmental state. Proc Natl Acad Sci U S A 107, 21931–21936. 1016071107.

49. Liu, Z., Zhang, Y., Ma, N., Yang, Y., Ma, Y., Wang, F., Wang, Y., Wei, J., Chen, H., Tartarone, A., et al. (2023). Progenitor-like exhausted SPRY1(+)CD8(+) T cells potentiate responsiveness to neoadjuvant PD-1 blockade in esophageal squamous cell carcinoma. Cancer Cell 41, 1852–1870 e1859. 10.1016/j.ccell.2023.09.011.

50. Deng, Q., Han, G., Puebla-Osorio, N., Ma, M.C.J., Strati, P., Chasen, B., Dai, E., Dang, M., Jain, N., Yang, H., et al. (2020). Characteristics of anti-CD19 CAR T cell infusion products associated with efficacy and toxicity in patients with large B cell lymphomas. Nat Med 26, 1878–1887. 10.1038/s41591-020-1061-7.

51. Haradhvala, N.J., Leick, M.B., Maurer, K., Gohil, S.H., Larson, R.C., Yao, N., Gallagher, K.M.E., Katsis, K., Frigault, M.J., Southard, J., et al. (2022). Distinct cellular dynamics associated with response to CAR-T therapy for refractory B cell lymphoma. Nat Med 28, 1848–1859. 10.1038/s41591-022-01959-0.

52. Chu, Y., Dai, E., Li, Y., Han, G., Pei, G., Ingram, D.R., Thakkar, K., Qin, J.-J., Dang, M., and Le, X. (2023). Pan-cancer T cell atlas links a cellular stress response state to immunotherapy resistance. Nature medicine 29, 1550–1562.

53. Baessler, A., and Vignali, D.A. (2024). T cell exhaustion. Annual review of immunology 42, 179–206.

54. Wang, F., Cheng, F., and Zheng, F. (2022). Stem cell like memory T cells: A new paradigm in cancer immunotherapy. Clinical Immunology 241, 109078.

55. Gattinoni, L., Klebanoff, C.A., and Restifo, N.P. (2012). Paths to stemness: building the ultimate antitumour T cell. Nature Reviews Cancer 12, 671–684.

56. Rade, M., Böhlen, S., Neuhaus, V., Löffler, D., Blumert, C., Merz, M., Köhl, U., Dehmel, S., Sewald, K., and Reiche, K. (2023). A time-resolved meta-analysis of consensus gene expression profiles during human T-cell activation. Genome biology 24, 287.

57. Smith-Garvin, J.E., Koretzky, G.A., and Jordan, M.S. (2009). T cell activation. Annual review of immunology 27, 591–619.

58. Neefjes, J., Jongsma, M.L., Paul, P., and Bakke, O. (2011). Towards a systems understanding of MHC class I and MHC class II antigen presentation. Nature reviews immunology 11, 823–836.

59. Pishesha, N., Harmand, T.J., and Ploegh, H.L. (2022). A guide to antigen processing and presentation. Nature Reviews Immunology 22, 751–764.

60. Golstein, P., and Griffiths, G.M. (2018). An early history of T cell-mediated cytotoxicity. Nature Reviews Immunology 18, 527–535.

61. Voskoboinik, I., Whisstock, J.C., and Trapani, J.A. (2015). Perforin and granzymes: function, dysfunction and human pathology. Nature Reviews Immunology 15, 388–400.

62. Platanias, L.C. (2005). Mechanisms of type-I-and type-II-interferon-mediated signalling. Nature Reviews Immunology 5, 375–386.

63. Ivashkiv, L.B., and Donlin, L.T. (2014). Regulation of type I interferon responses. Nature Reviews Immunology 14, 36–49.

64. Steffin, D., Ghatwai, N., Montalbano, A., Rathi, P., Courtney, A.N., Arnett, A.B., Fleurence, J., Sweidan, R., Wang, T., Zhang, H., et al. (2025). Interleukin-15-armoured GPC3 CAR T cells for patients with solid cancers. Nature 637, 940–946. 10.1038/s41586-024-08261-8.

65. Goswami, S., Pauken, K.E., Wang, L., and Sharma, P. (2024). Next-generation combination approaches for immune checkpoint therapy. Nat Immunol 25, 2186–2199. 10.1038/s41590-024-02015-4.

66. Wang, S.L., and Chan, T.A. (2025). Navigating established and emerging biomarkers for immune checkpoint inhibitor therapy. Cancer Cell 43, 641–664. 10.1016/j.ccell.2025.03.006.

67. Pan, X., Zhang, Y.W., Dai, C., Zhang, J., Zhang, M., and Chen, X. (2025). Applications of mRNA Delivery in Cancer Immunotherapy. Int J Nanomedicine 20, 3339–3361. 10.2147/IJN.S500520.

68. Khan, S.H., Choi, Y., Veena, M., Lee, J.K., and Shin, D.S. (2024). Advances in CAR T cell therapy: antigen selection, modifications, and current trials for solid tumors. Front Immunol 15, 1489827. 10.3389/fimmu.2024.1489827.

69. Singh, N., Frey, N.V., Engels, B., Barrett, D.M., Shestova, O., Ravikumar, P., Cummins, K.D., Lee, Y.G., Pajarillo, R., Chun, I., et al. (2021). Antigen-independent activation enhances the efficacy of 4-1BB-costimulated CD22 CAR T cells. Nat Med 27, 842–850. 10.1038/s41591-021-01326-5.

70. Qi, C., Liu, D., Liu, C., Wei, X., Ma, M., Lu, X., Tao, M., Zhang, C., Wang, X., He, T., et al. (2024). Antigen-independent activation is critical for the durable antitumor effect of GUCY2C-targeted CAR-T cells. J Immunother Cancer 12. 10.1136/jitc-2024-009960.

71. Soni, U.K., Chadchan, S.B., Gupta, R.K., Kumar, V., and Kumar Jha, R. (2021). miRNA-149 targets PARP-2 in endometrial epithelial and stromal cells to regulate the trophoblast attachment process. Mol Hum Reprod 27. 10.1093/molehr/gaab039.

72. Ponder, J.W., and Case, D.A. (2003). Force fields for protein simulations. Advances in protein chemistry 66, 27–85.

73. Rohl, C.A., Strauss, C.E., Misura, K.M., and Baker, D. (2004). Protein structure prediction using Rosetta. In Methods in enzymology, (Elsevier), pp. 66–93.

74. Abraham, M.J., Murtola, T., Schulz, R., Páll, S., Smith, J.C., Hess, B., and Lindahl, E. (2015). GROMACS: High performance molecular simulations through multi-level parallelism from laptops to supercomputers. SoftwareX 1, 19–25.

75. Evans, D.J., and Holian, B. (1985). The nose–hoover thermostat. A)’=(A Is) 1, 18.

76. Parrinello, M., and Rahman, A. (1981). Polymorphic transitions in single crystals: A new molecular dynamics method. Journal of Applied physics 52, 7182–7190.

77. Hess, B., Bekker, H., Berendsen, H.J., and Fraaije, J.G. (1997). LINCS: A linear constraint solver for molecular simulations. Journal of computational chemistry 18, 1463–1472.

